# Discovery of nanobodies against SARS-CoV-2 and an uncommon neutralizing mechanism

**DOI:** 10.1101/2021.07.20.453054

**Authors:** Tingting Li, Bingjie Zhou, Zhipu Luo, Yanling Lai, Suqiong Huang, Yuanze Zhou, Anupriya Gautam, Salome Bourgeau, Shurui Wang, Juan Bao, Jingquan Tan, Dimitri Lavillette, Dianfan Li

**Author notes:** Equal contribution (Tingting Li, Bingjie Zhou, Zhipu Luo).

## Abstract

SARS-CoV-2 and its variants continue to threaten public health. The virus recognizes the host cell by attaching its Spike receptor-binding domain (RBD) to the host receptor ACE2. Therefore, RBD is a primary target for neutralizing antibodies and vaccines. Here we report the isolation, and biological and structural characterization of two single-chain antibodies (nanobodies, DL4 and DL28) from RBD-immunized alpaca. Both nanobodies bind Spike with affinities that exceeded the detection limit (picomolar) of the biolayer interferometry assay and neutralize the original SARS-CoV- 2 strain with IC_50_ of 0.086 μg mL^-1^ (DL4) and 0.385 μg mL^-1^ (DL28). DL4 and a more potent, rationally designed mutant, neutralizes the Alpha variant as potently as the original strain but only displays marginal activity against the Beta variant. By contrast, the neutralizing activity of DL28, when in the Fc-fused divalent form, was less affected by the mutations in the Beta variant (IC_50_ of 0.414 μg mL^-1^ for Alpha, 1.060 μg mL^-1^ for Beta). Crystal structure studies reveal that DL4 blocks ACE2-binding by direct competition, while DL28 neutralizes SARS-CoV-2 by an uncommon mechanism through which DL28 distorts the receptor-binding motif in RBD and hence prevents ACE2-binding. Our work provides two neutralizing nanobodies for potential therapeutic development and reveals an uncommon mechanism to design and screen novel neutralizing antibodies against SARS-CoV-2.

## INTRODUCTION

A key step for SARS-CoV-2 infection is the molecular engagement between the receptor-binding domain (RBD) on the Spike protein (S) and the human receptor angiotensin-converting enzyme 2 (ACE2) ^1–4^. S is a heavily glycosylated trimeric protein that in the pre-form contains 1273 amino acid residues. Upon cleavage by host proteases, S breaks down to two subunits S1 and S2 at a region near residue 685. RBD (residues 330-526) is contained in the S1 subunit ^4^. In the pre-fusion state, S exists in multiple conformations regarding the relative position of RBD to the rest of the protein. In its ‘closed’ conformation, all three subunits are very similar and the receptor-binding motif (RBM) of the RBD is buried by adjacent N-terminal domains (NTDs) of S1. The RBD in the closed S is referred to as the ‘down’ conformation and they are incompetent to engage with ACE2. In the ‘open’ state, one, two, or all three RBDs could assume the ‘up’ conformation, exposing the RBM to engage with ACE2 ^1, 2, 5, 6^. Reflecting the importance of ACE2-RBD binding in viral infection, hundreds of existing neutralizing antibodies target this event by direct blockage, steric hindrance, or locking the RBDs in the ‘down’ conformation ^7^.

The single-chain camelids-derived antibodies possess attractive features ^8^. The variable region of the heavy-chain antibodies is referred to as nanobodies owing to their small sizes (∼14 kDa). Despite having a single chain, nanobodies can target antigens with comparable selectivity and affinity to conventional antibodies. Being small, nanobodies are ultra-stable, relatively easy to produce (in microbial) with low costs and high yields, and amenable to protein engineering such as fusion in various forms. Such fusion can result in improved potency – binding affinity and neutralizing activity can increase by hundreds to thousands of fold ^9–11^. In addition, nanobodies that recognize non-competing epitopes can be conveniently fused to make biparatopic nanobodies that are potentially more tolerant to escape mutant strains ^9, 10, 12^. The heat stability of nanobodies opens the possibility of using them as inhaling drugs for respiratory diseases ^8^ (and indeed potentially for SARS-CoV-2 as demonstrated in hamsters ^13^) and offers convenience in storage and transport. In the past months, dozens of neutralizing nanobodies against SARS-CoV-2 have been reported ^10–21^.

A challenge in developing neutralizing antibodies and vaccines against viruses is their ability to mutate. In particular, mutations in RBD that retain its structural integrity and function (ACE2-binding) may escape neutralizing antibodies by altering the binding surface either in composition or in conformation, or both ^22–25^. In the past months, strains such as the lineage B1.1.7 and B1.351, referred to as the UK and South African (SA) variant based on the region they first emerge, or the Alpha and Beta strain by the recent recommendation from the World Health Organization, have caused outbreaks and concerns about how they could change the course of the pandemic because their high virulence and their general resistance against antibodies and vaccines that were developed using previous strains ^26, 27^. Indeed, the Delta strain which is the prevalent strain in the recent global outbreaks ^28^ contains both the mutations seen in UK and SA variants although it is yet to be established how much this ‘double mutant’ compromises the protective effect of the current vaccines. Of relevance, a laboratory-generated mutant, E406W ^29^, could escape a Regeneron cocktail that contains two mAbs recognizing different epitopes on RBD. Given the large number of active cases, it is reasonable to assume that more escape mutants are almost certain to emerge. Due to the lag phase between outbreaks caused by new mutants and the development of vaccines/mAbs against the mutants, it is of vital importance to have different antibodies and to test and develop strategies to identify antibodies with broad reactivity.

Here, we report the selection and structural characterization of two RBD-targeting neutralizing nanobodies (dubbed DL4 and DL28) isolated from immunized alpaca. DL4 binds the Spike tightly at the RBM with a *K*_D_ below the detection limit of our biolayer interferometry assay. DL4 neutralizes the Alpha but not the Beta variant. By contrast, DL28 recognizes RBD at a region adjacent to RBM and is less affected by the mutations in the Beta variant. Structural characterizations rationalize their variable potency against different SARS-CoV-2 strains and suggest an unreported neutralization mechanism by which DL28 distorts the RBM and diminishes ACE2-binding. Our work adds more evidence that RBM-antibodies are more prone to escape mutants and identifies nanobodies and its associated epitope for therapeutic development against SARS-CoV-2 mutants.

## RESULTS

### Isolation of a high-affinity neutralizing nanobody from immunized alpaca

An adult female alpaca was immunized four times using recombinantly expressed RBD. ELISA test of sera showed an antibody titer of ∼1 × 10^6^ after four rounds of immunization compared with the pre-immunization sample. mRNA isolated from peripheral blood lymphocytes of RBD-immunized alpaca was reverse-transcripted into cDNA for the construction of a phage display library (**Fig. 1A**). Three rounds of solution panning were performed with increasingly stringent conditions and an off-selection step to screen high-affinity nanobodies. Subsequence screening using ELISA and fluorescence-detection size exclusion chromatography (FSEC) ^10^ identified binders with ELISA signal that is at least 3 times higher than a control nanobody, as well as the ability to shift the gel filtration peak of fluorescently labeled RBD at 0.5 μM (**Fig. 1A**). We identified 28 unique clones as positive clones, among which DL4 was first chosen based on its ability to cause earlier elution of RBD in FSEC (**Fig. 1B**) and its exceptional binding kinetics (**Fig. 1C**). The binding affinity between DL4 and S exceeded the detection limit for the biolayer interferometry assay on an Octet system, reporting a *K*_D_ of <1 pM and a slow *k*_off_ of <1.0 × 10^-7^ s^-1^ (**Fig. 1C**). A neutralization assay using SARS-CoV-2 pseudotyped particles (pp) bearing the S from the original Wuhan strain displayed an IC_50_ of 0.086 μg mL^-1^ for DL4 (**Fig. 1D**). **Table S1** summarizes the sequence and neutralizing activity of DL4 and all other nanobodies in this study.

**Fig. 1.**
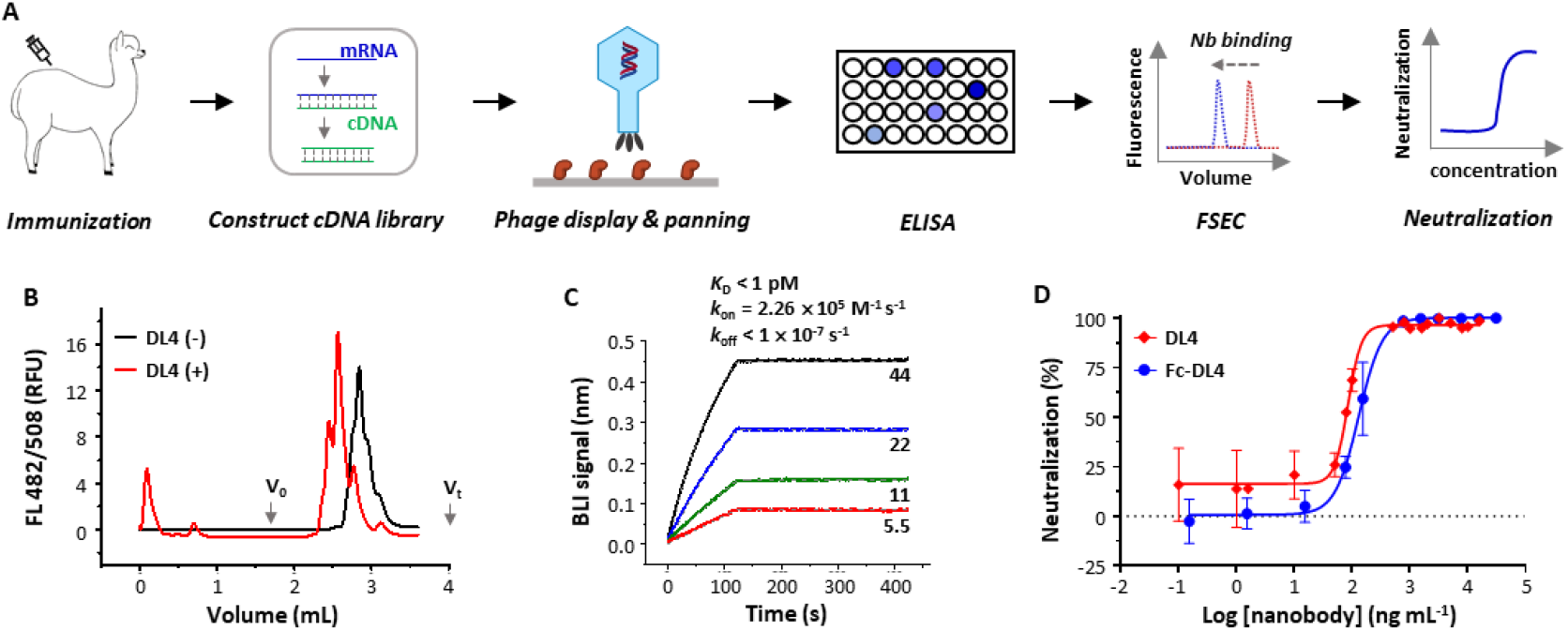
Strategy and isolation of neutralizing nanobodies. (**A**) Schematic flowchart for identification of neutralizing nanobodies (Nbs). mRNA was isolated from an alpaca that was immunized with the receptor-binding domain. A phage display library expressing nanobodies was selected against RBD. Positive clones were screened using enzyme-linked immunosorbent assay (ELISA) and fluorescence-detector size exclusion chromatography (FSEC) for RBD-binding, and purified nanobodies were screened using neutralization assays with SARS-CoV-2 pseudoviruses. (**B**) Unpurified DL4 causes earlier elution of the fluorescently labeled RBD on analytic gel filtration. (**C**) Binding kinetics of DL4 to Spike using biolayer interferometry (BLI) with DL4 immobilized and Spike as analyte at indicated concentrations (nM). Solid lines indicate original data and dotted lines (grey) indicate fitted curve. (**D**) Neutralization assay of DL4 and Fc-DL4 against SARS-CoV-2 pseudoviruses. Error bar presents s.d. from three independent experiments.

### Structural characterization of the DL4 epitope

To accurately characterize the epitope of DL4, we crystallized the DL4-RBD complex in the space group of *P*22_1_2_1_ and solved its structure to 1.93 Å resolution by molecular replacement using published RBD and nanobody structures as search models. The structure was refined to *R*_work_ / *R*_free_ of 0.1973 / 0.2351 with no geometry violations (**Table S2**). Each asymmetric unit contains two DL4-RBD complexes which are highly similar with Cα RMSD of 0.207 Å. Chains A and B are used for structure description.

The RBD structure assembles a high-chair shape and DL4 binds RBD at the ‘seat’ and ‘backrest’ region with a buried surface area ^30^ of 957.6 Å^2^ (**Fig. 2A**), with contributions of 151.0 Å^2^ from CDR1, 322.6 Å^2^ from CDR2, 240.5 Å^2^ from CDR3, and interestingly, 243.5 Å^2^ (25 % of the total surface) from the framework region. For clarity, we label residues from RBD with a prime. The three CDRs interact with RBD via two salt bridge pairs (Glu30 / Arg403’ and Arg50 / Glu484’), four hydrogen bonds (Thr33 / Gln493’, Asn54 / Asn450’, Gln101 / Leu 455’), cation-π interactions (Arg50 / Phe 490’), and hydrophobic interactions by apolar residues or hydrocarbon potion of polar residues such as Glu484’. The framework loop contributed a hydrogen bond (Asp74 / Gly446’) and a cation-π interaction (Arg71 / Tyr449’) (**Fig. 2B-3D**).

**Fig. 2.**
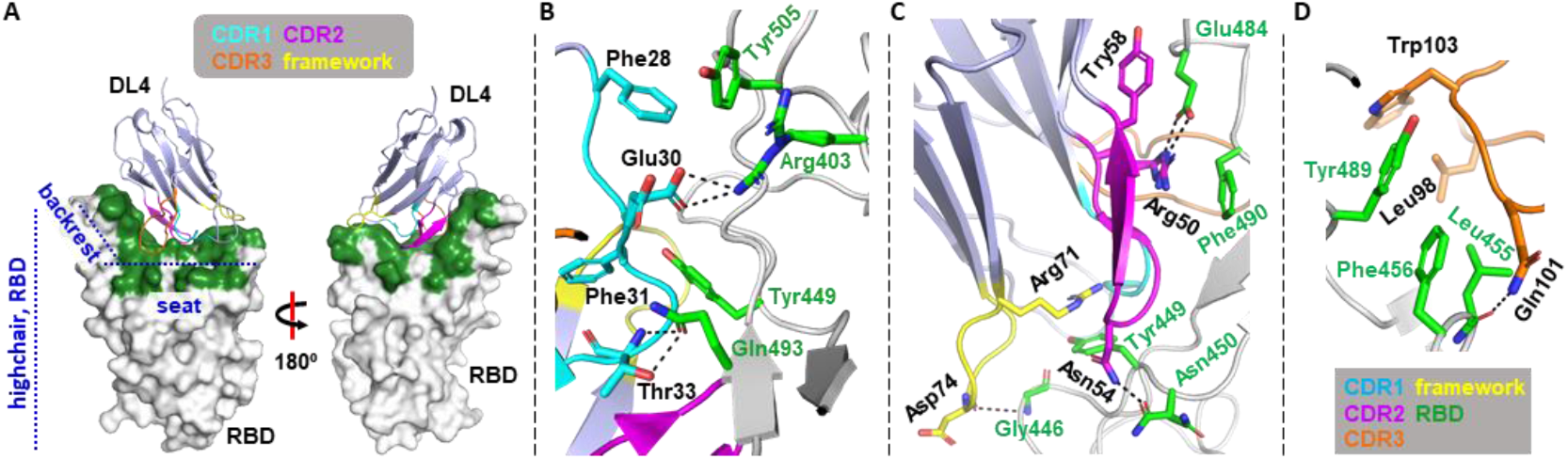
Crystal structure of the DL4 in complex with the receptor-binding domain (RBD). (**A**) The overall structure of the DL4 (light blue) in complex with RBD (white). DL4 binds the highchair-shaped RBD at the ‘seat’ and ‘backrest’ region. The binding interface is colored green. Three CDRs and the framework residues involved in the binding are color-coded as indicated. (**B-D**) Detained interaction between DL4 and RBD. **B**, CDR1; **C**, CDR2 and framework; **D**, CDR3. Dash lines indicate distances within 3.8 Å. Sidechain of the RBD residues are colored green.

**Fig. 3.**
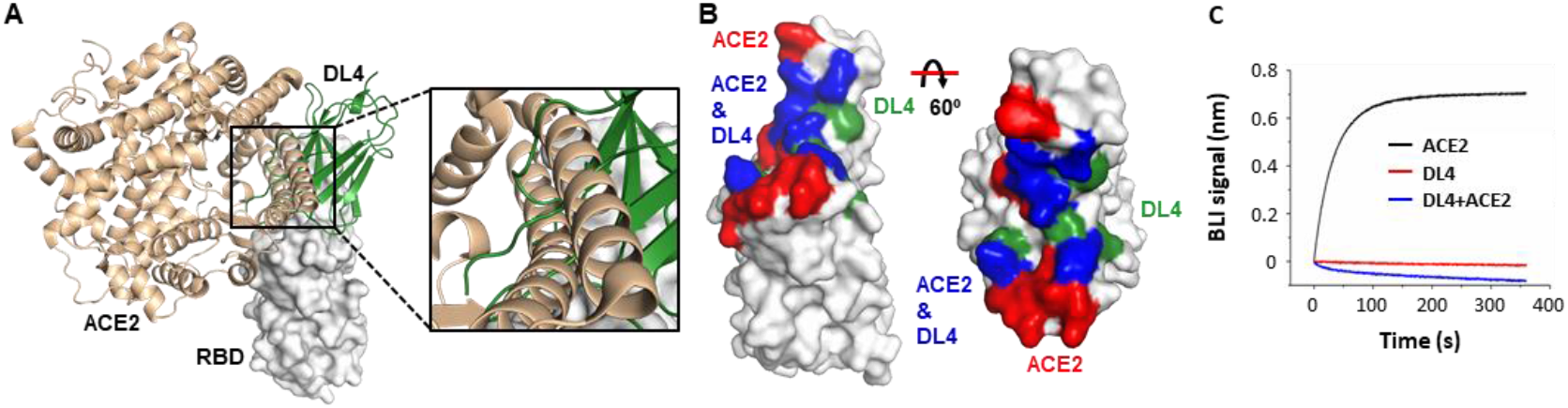
Nanobody DL4 engages the receptor-binding domain (RBD) at the receptor-binding motif and directly competes with ACE2 for RBD-binding. (**A**) Aligning the DL4-RBD structure onto the ACE2-RBD structure (PDB ID 6M0J) ^31^ reveals clashes between ACE2 (wheat) and DL4 (green). Only the RBD from the DL4- binding structure is shown (white). (**B**) The overlap (blue) between the ACE2-binding site (red) and the DL4 epitope (green). (**C**) Pre-incubation of DL4 with RBD prevents ACE2 from binding to RBD. A sensor coated with RBD was first saturated with DL4 before incubated with a DL4-containing solution with (blue) or without (red) ACE2. As a control, the ACE2-RBD binding profile (black) was recorded using the same procedure without DL4 on a biolayer interferometry (BLI) system.

### DL4 competes directly with ACE2 for RBD-binding

Aligning the DL4-RBD complex to the ACE2-RBD structure ^31, 32^ reveals a large overlap between the DL4 epitope and the receptor-binding motif (RBM) (**Fig. 3A, 4B**). Specifically, the shared site includes 15 residues, some of which, such as Gln493’ and Glu484’ are key residues for both the receptor- and DL4-binding. Consistent with the structural observation, pre-incubation with DL4 completely blocked the binding between ACE2 and RBD (**Fig. 3C**).

**Fig. 4.**
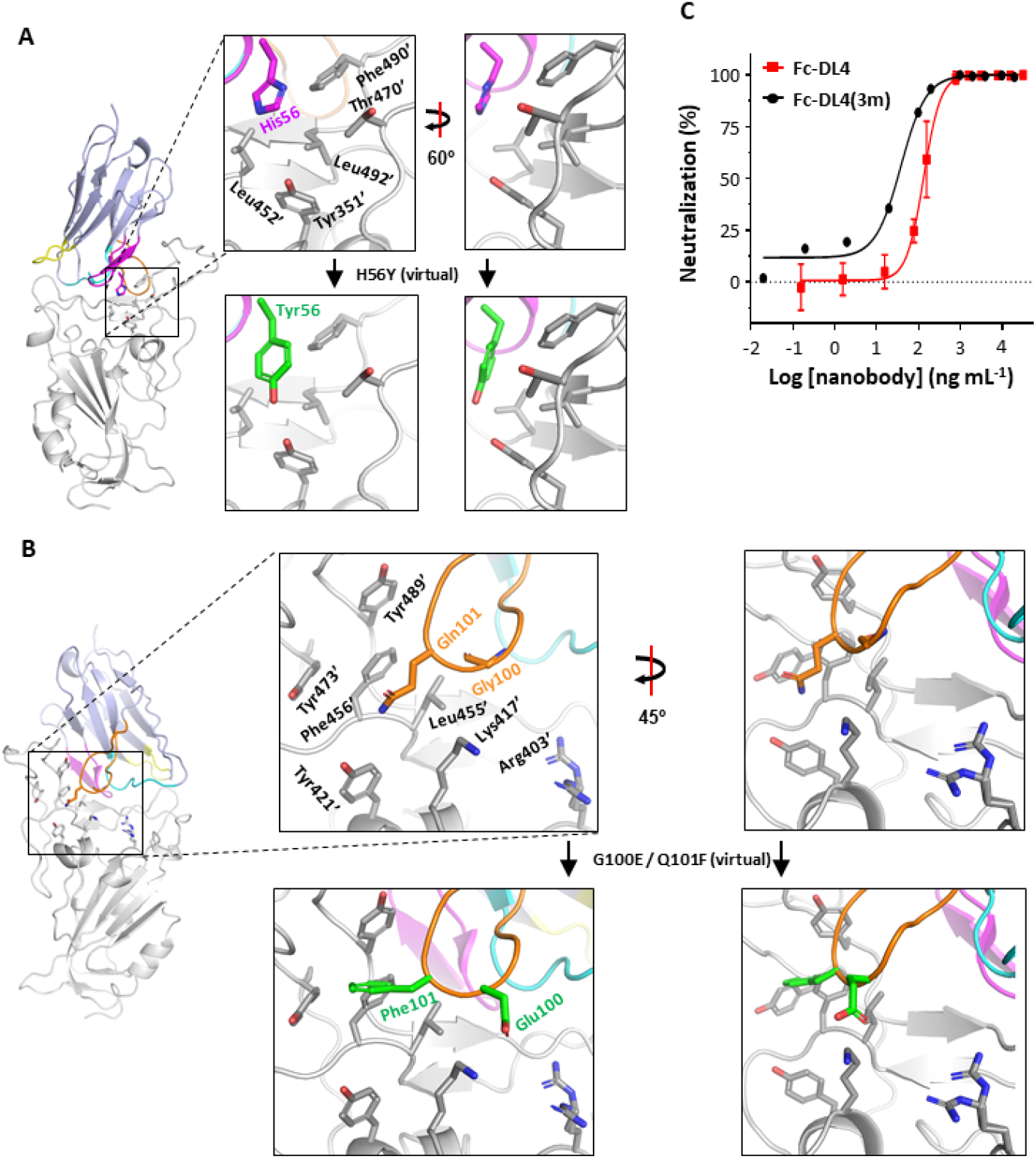
Structure-based design for a gain-of-function DL4 mutant. (**A, B**) The rationale for the design of H56Y (**A**) and G100E and Q101F (**B**). H56Y and Q101F may bind RBD tighter because of hydrophobic matching. G100E may bind RBD tighter by introducing a salt bridge. (**C**) The triple mutant displayed a 4-fold neutralizing activity compared to the wild-type DL4. Data for Fc-DL4 are from Fig. 1D. Data for Fc-DL4 are the average from two independent experiments.

The DL4 structure was also aligned to the S to assess RBD-binding in the context of the trimer structure. As shown in **Fig. S1**, the DL4-epitope is well exposed and no clashes are observed for DL4 on the ‘up’-RBD. Owing to its small size, only minor clashes are apparent when DL4 is aligned to the ‘down’-RBDs in both the open and closed conformations. The clashes are contributed by the Asn343-linked glycans and nearby residues. The structural analysis suggests the DL4 epitope in S is accessible, in accord with the BLI binding results (**Fig. 1C**).

### Structure-based design improved DL4’s potency

Next, we set to engineer DL4 for higher neutralizing activity. Avidity effects are commonly exploited for nanobody engineering ^10, 33^ and we also constructed the Fc version of DL4. Unlike those in previous reports, however, the Fc fusion did not increase neutralizing activity, displaying an IC_50_ of 0.132 μg mL^-1^, which, by molarity (3.4 nM), was similar to that of the monomeric DL4 (5.3 nM).

Previously, we have designed gain-of-function nanobody mutations based on structural information to increase binding affinity and neutralizing activity ^10^. This approach was used again for DL4. Analyzing the DL4-RBD structure reveals that His56 from CDR2 is located in a hydrophobic microenvironment (**Fig. 4A**) and does not contribute to hydrogen bonding (**Fig. 2C**). To match the hydrophobic patch, His56 was mutated to Phe, Tyr, and Trp. Similarly, Gln101 in CDR3 was also mutated to the three aromatic residues to match the hydrophobic patch on the RBD made by Tyr421’, Leu455’, Phe456’, Try473’, Tyr489’, and the hydrocarbon portion of Lys471’(**Fig. 4B**). In addition, the G100E mutant was designed to introduce a possible salt bridge with Lys417’ or the nearby Arg403’. Subsequent neutralizing assays identified H56Y, Q101F, and G100E as gain-of-function mutants with IC_50_ values of 0.133, 0.098, and 0.084 μg mL^-1^, respectively (the Fc-version was used, **Fig. S2A**). Consistently, the triple mutant showed a 3-fold increase of neutralizing activity, with an IC_50_ of 0.038 μg mL^-1^ (0.49 nM) (**Fig. 4C**).

### DL4 neutralizes the Alpha potently but neutralizes the Beta variant poorly

A challenge in antibody and vaccine research against SARS-CoV-2 is its ability to evolve escape mutants. During the study, two major more infectious variants, the lineages B1.1.7 (Alpha) and B1.353 (Beta), were reported. The Alpha strain contains the N501Y mutation in the RBD and the Beta strain contains two additional mutations, K417N and E484K.

Although Ans501’ is in the vicinity of the CDR1, it does not form hydrogen bonds with DL4 (**Fig. 2B**). Therefore, mutation of Ans501’ is not expected to affect DL4-RBD binding, at least directly. In addition, a tyrosine replacement appeared to be compatible with the local hydrophobic patch consisting of Phe28/29/31; and Tyr501’ may even form a hydrogen bond with Glu30 (**Fig. 5A**). Therefore, it was expected that DL4 should remain competent against Alpha. This was indeed the case for both DL4 and the triple mutant H56Y/G100E/Q101F; they displayed equal or slightly higher neutralizing activity against Alpha compared to the original Wuhan strain (**Fig. 5B**).

**Fig. 5.**
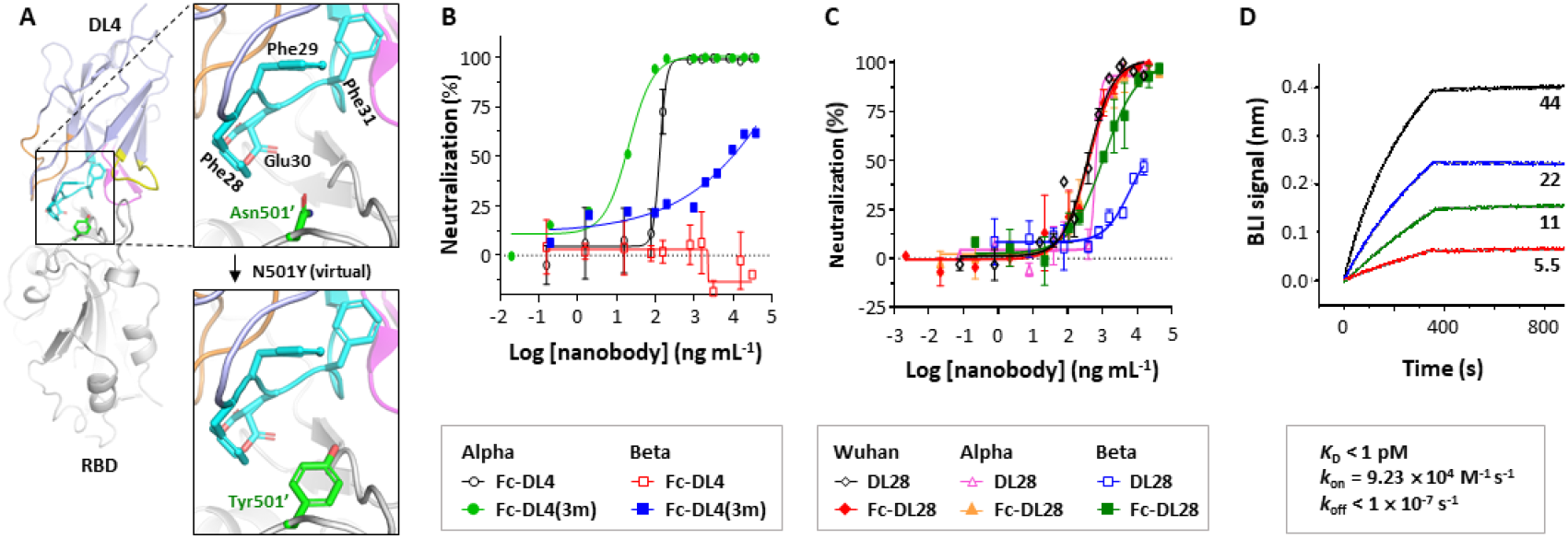
Varying efficiency of nanobodies DL4 and DL28 in neutralizing the original SARS-CoV-2 and the Alpha and the Beta variants. (**A**) A possible structural reason for DL4’s potent neutralizing activity against the Alpha strain. (**B**) DL4 neutralizes the Alpha variant but not the Beta variant. Data for Fc-DL4(3m) are the average from two independent experiments. (**C**) DL28 neutralizes the original Wuhan strain, and the Alpha and Beta variants. Error bar presents s.d. from three independent experiments. (**C**) Binding kinetics of DL28 to Spike using biolayer interferometry (BLI) with DL28 immobilized and Spike as analyte at indicated concentrations (nM). Solid lines indicate original data and dotted lines (grey) indicate fitted curve. In **B** and **C**, error bars represent the standard error (s.d.) for data from three independent experiments.

The Beta variant contains a lysine replacement of Glu484’, a residue that forms a key salt bridge with Arg50 in CDR2. The E484K mutation would not only eliminate the salt bridge, but also introduce charge-charge repulsion with Arg50. Thus, the Beta variant was expected to escape DL4. This was also confirmed by the neutralizing assay using both Fc-DL4 and the triple mutant (**Fig. 5B**). Interestingly, two DL4 mutants (R50E and R50D), designed to restore the salt bridges with Lys484’ in the Beta variant RBD, could not neutralize the Beta variant.

### Identification of a nanobody capable of neutralizing the Beta variant

One of the focuses in the research of SARS-CoV-2 neutralizing antibodies is to identify antibodies with broad reactivity. To this end, we re-screened clones and obtained a nanobody (dubbed DL28) that showed weak neutralizing activity against the Beta variant. Increasing avidity by Fc-fusion improved the neutralizing activity, with an IC_50_ of 1.06 μg mL^-1^ which is ∼2 fold of that for the original Wuhan strain (**Fig. 5C**). Similar to DL4, DL28 could bind to the S protein with ultra-high affinity – its *K*_D_ was also below the detection limit of the BLI assay (**Fig. 5D**).

### Structural characterization of the DL28 epitope

To characterize the epitope, we also crystallized the DL28-RBD complex. The crystals belong to the space group of *P*6_5_22 and diffracted to 3.0 Å at the synchrotron. The structure was refined to *R*_work_ / *R*_free_ of 0.2264 / 0.2467 with no geometry violations (**Table S2**). The asymmetric unit contains two copies of complexes that are very similar (Cα RMSD of 0.509 Å). The chains A/C are used for structure description.

DL28 binds RBD at one side of the high-chair shape with a buried surface area of 986.3 Å^2^ (**Fig. 6A**). In addition to three CDRs (CDR1, 41.9 Å^2^; CDR2, 195.4 Å^2^; CDR3, 369.2 Å^2^), the framework region also contributed significantly to the binding with a buried surface area of 379.8 Å^2^ (∼40% of the total). Characteristically, most interactions are contained in CDR3 and only one residue in CDR1 is involved in the binding (**Fig. 6B-6E**). Overall, the interaction involves 12 hydrogen bonds and a π-π interaction between Phe47 and Phe450’.

**Fig. 6.**
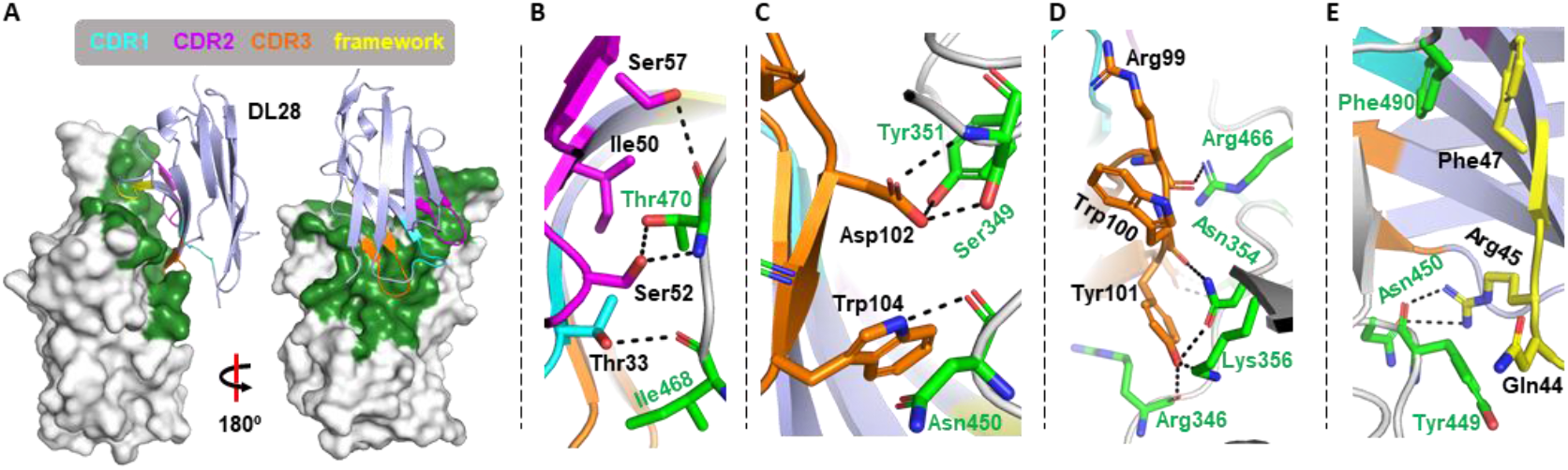
Crystal structure of the DL28 in complex with the receptor-binding domain (RBD). (**A**) The overall structure of the DL4 (light blue) in complex with RBD (white). DL4 binds the highchair-shaped RBD at one side. The binding interface is colored green. Three CDRs and the framework residues involved in the binding are color-coded as indicated. (**B-D**) Detained interaction between DL4 and RBD. **B**, CDR1; **C**, CDR2 and framework; **D**, CDR3. Dash lines indicate distances within 3.8 Å. The sidechain of the RBD residues are colored green.

Similar to DL4, aligning the DL28 structure to the S structure reveals no clashes for DL28 in binding with the ‘up’-RBD, and only minor clashes with the NTD from the clock-wise subunit when binding with the ‘down’-RBD (**Fig. S3**).

### DL28 impairs ACE2-binding mainly by RBM distortion

Cross-competition binding assays showed that DL28 also blocked receptor binding to near completion (**Fig. 7A**). Aligning the DL28-RBD structure to the ACE2- RBD ^31^ revealed that DL28 and ACE2 approach RBD at the opposite sides of the ‘seat’ region. Unlike DL4, only minor clashes were observed between aligned DL28 and ACE2. Specifically, Gln44 of DL28 would clash with the mainchain of the ACE2 α- helix α_20-52_ (subscript numbers refer to the start-end residues) (**Fig. 7B**), which contains most of the key receptor-RBD interactions ^31^. Mutating Gln44 to glycine resulted in a slight increase in neutralizing activity (**Fig. S4, Table S2**). Further mutation of the adjacent Lys43 to glycine resulted in a somewhat weaker activity for ACE2-blocking (**Fig. 7C**) and neutralization (**Fig. S4, Table S2**). Possibly, the tri-glycine motif (together with Gly42) introduces structural instability to the nanobody framework and affects the orientation of the CDRs for tight binding. The results suggest that the steric hindrance is not the main factor for neutralization.

**Fig. 7.**
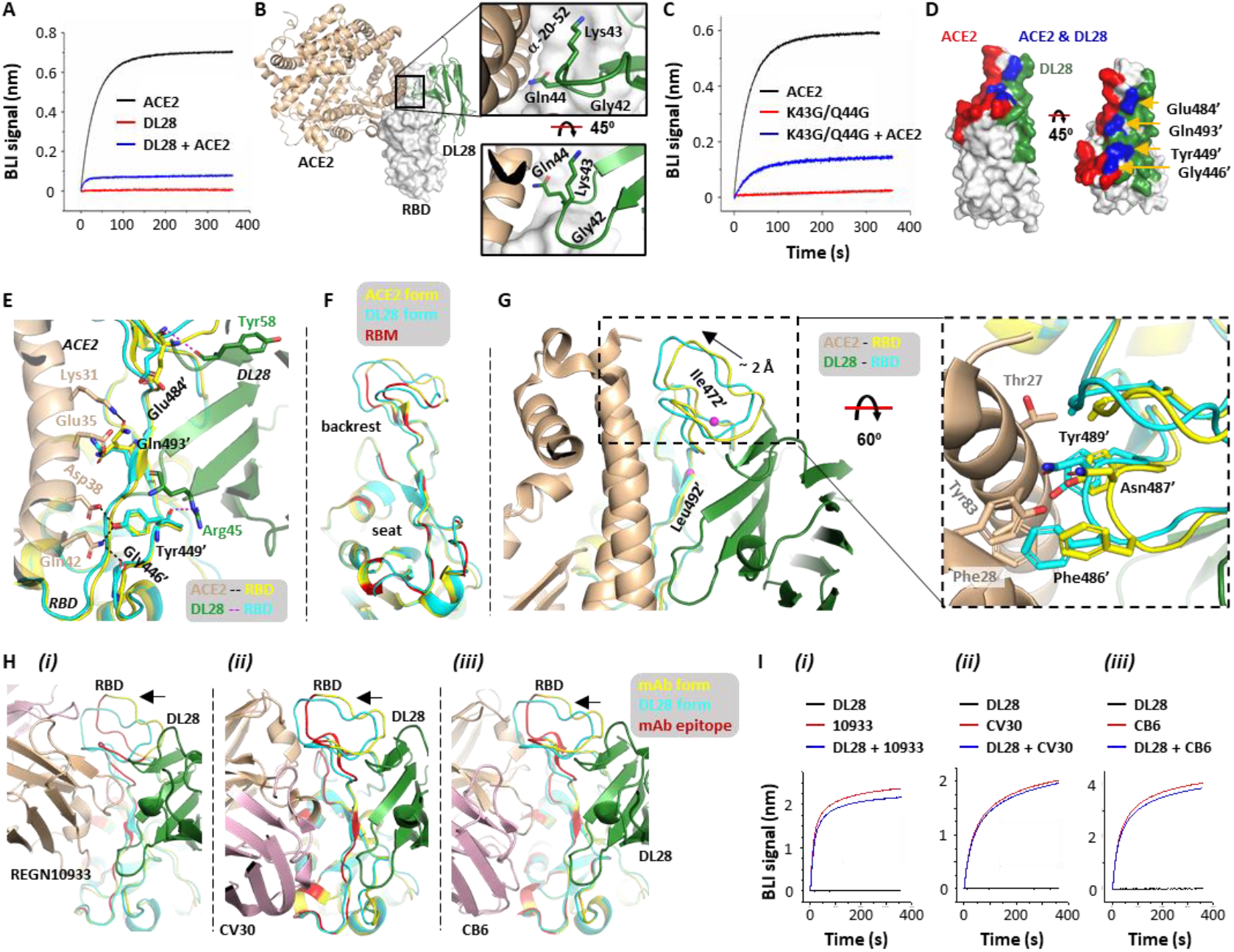
The distortion of the receptor-binding motif (RBM) by DL28 affects RBD-binding for ACE2 but not for three RBM-targeting monoclonal antibodies. (**A**) Pre-incubation of DL28 with RBD inhibits ACE2 for RBD-binding. A sensor coated with RBD was first saturated with DL28 before incubated with a DL28-containing solution with (blue) or without (red) ACE2. As a control, the ACE2-RBD binding profile (black) was recorded using the same procedure without DL28 on a biolayer interferometry (BLI) system. (**B, C**) The minor clashes between DL28 and ACE2 do not play a major role in cross-competition. (**B**) Gln44 on DL28 is in close contact with the RBD-interacting α-helix from ACE2 when the DL28-RBD structure is aligned onto the ACE2-RBD structure. (**C**) The triple-glycine DL28 (Gly42, K43G/Q44G) retained the ability to inhibit ACE2 for RBD-binding. The experimental setting was the same as in A. (**D, E**) The epitope of DL28 (green) overlaps with the ACE2-binding site (RBM, red) (**D**), but the overlap does not appear to mutually affect their binding with RBD (**E**). Glu484’, Gln493’, and Tyr449’ all interact with ACE2 (wheat) with sidechains and interact with DL28 (green) with mainchain atoms. Black dashed lines indicate the interactions between ACE2 and RBD (yellow). Magenta dashed lines indicate the interactions between DL28 and RBD (cyan). (**F**) Comparison of the RBD conformations at the RBM between the ACE2-bound (yellow) ^31^ and the DL28-bound (cyan) forms. The ACE2-interacting residues are colored red. The distorted part (backrest) and the non-distorted part (seat) are marked. (**G**) Alignment of the DL28- RBD structure (green and cyan) with the ACE2-RBD structure ^31^ (wheat and yellow). The RBM distortion caused by DL28 binding leads to clashes between the ‘backrest’ part of the RBM and the two RBD-interacting α-helices in ACE2 (wheat). (**H**) Alignment of the DL28-RBD structure (green and cyan) ^31^ with the structures of REGN10933-RBD ^34^ (wheat/pink and yellow, *i*), CV30-RBD ^35^ (*ii*), and CB6-RBD ^36^ (*iii*). (**I**) DL28 forms non-competing pairs with REGN10933 (*i*), CV30 (*ii*), and CB6 (*iii*) for RBD-binding. A sensor coated with RBD was first saturated with DL28 before incubated with a DL28-containing solution with (blue) or without (red) other antibodies. As a control, the binding between RBD and other antibodies (black) was recorded using the same procedure without DL4 on a biolayer interferometry (BLI) system.

Mapping the DL28 epitope and the RBM to the RBD reveals that they overlap by four residues, namely Gly446’, Tyr449’, Glu484’, and Gln493’ (**Fig. 7C**). However, none of the four RBD residues interacts with DL28 by sidechain (Gly446’ has no sidechain). Gly446’ and Gln493’ are only in the proximity without specific hydrogen bonds with DL28; and although Tyr449’ and Glu484’ form hydrogen bonds with Arg45 and Tyr58, the interactions only involved mainchain of Tyr449’/Gln493’. In addition, both the mainchain and sidechain of the four RBD residues showed negligible differences in conformation between the ACE2- and DL28-bound forms (**Fig. 7D**). Therefore, the RBD conformation at this overlapping region appears to be compatible for simultaneous binding with ACE2 and DL28. The analysis also supports the abovementioned idea of additional mechanisms for DL28’s receptor-blocking activity (**Fig. 7A**).

Aligning the RBD structures in the receptor- and DL28-binding mode reveals that the backbone of the RBM is largely similar but displays noticeable distortion at the ‘backrest’ region (**Fig. 4A**) between residues Ile472’ and Leu492’ (**Fig. 7E, 7F**). Specifically, DL28 nudges this loop toward the direction of ACE2 by ∼ 2 Å. Notably, the pushing by DL28 was not mediated by sidechains or loop regions which may tolerate clashes by assuming alternative conformations. Rather, it was mediated by a 4- residue β-sheet (β_56-59_) which is part of the stable nanobody framework made of four stacking β-sheets (**Fig. 7E**). The tight binding (**Fig. 5D**) and the rigidness of the nanobody core should therefore force and lock the loop in the left-ward position. As a consequence, the distorted loop clashes with the α_20-52_ of ACE2. Specifically, Phe486’,

Ans487’, and Tyr489’ from RBD would come into close contact with residues Thr27 and Phe28 in the ACE2 α_20-52_ and Tyr83 in an adjacent helix. Therefore, the conformational change in the ‘backrest’ loop appears to be incompatible with ACE2- binding, unless ACE2 can adapt to the conformational change, which, as reasoned below, would be unlikely.

The ACE2 α_20-52_ lies on top of RBD like a lever. The C-terminal half of the helix binds the ‘seat’ region of the RBD, and the N-terminal end binds with the ‘backrest’ region. A 2-Å distortion at the ‘backrest’ area acts like forces pushing the lever at one end (**Fig. 7G**). The α-helix would have to deform/break to adapt to such a dramatic distortion. However, α-helices are generally rigid and α_20-52_ contains no helix-destabilizing residues such as proline and glycine. Therefore, we propose that DL28 neutralizes SARS-CoV-2 by an ‘RBM distortion’ mechanism.

### RBM distortion does not affect the binding of several RBM-targeting antibodies

It might be expected that the RBM-targeting monoclonal antibodies (mAbs) are incompatible with DL28 because of DL28’s ability to distort RBM. However, this was not the case for three such mAbs (that are available to us): REGN10933 ^34^, CV30 ^35^, and CB6 ^36^. Thus, although their epitopes would also be shifted by a similar or more extent compared to the RBM of ACE2 (**Fig. 7H**), the mAbs bound to RBD in the presence of DL28 (**Fig. 7I**).

Unlike ACE2, mAbs bind to RBD with CDRs which are usually made of, or connected to the rigid framework by, flexible loops. This may have allowed the mAbs to adapt and to remain bound with RBD. Thus, the results are seemingly contradictory to the ACE2 competition but can be rationalized by the structural analysis.

## DISCUSSION

In this study, we report two high-affinity RBD binders isolated from immunized alpaca and their structural and biological characterization. Most monovalent RBD-targeting nanobodies bind S or RBD with *K*_D_ in the nanomolar ranges ^9-18, 21^. Monovalent nanobodies with *K*_D_ values in the low picomolar ranges include two RBM- type binders: Nb20 (10.4 pM) ^21^ from immunized llama and Nanosota-1C (157 pM) by in vitro maturation of a binder from a naïve llama/alpaca library ^37^. Remarkably, both DL4 and DL28 binds S with sub-picomolar affinities. This reinforces the notion that, despite their small sizes, nanobodies can bind antigens with comparable affinity with Fab which is four times in size. One of the reasons, as revealed in this study and previous structural reports, is that the framework region of the nanobodies can also participate in the antigen-binding, thus essentially expanding the binding surface and increasing the number of interactions. In addition, as revealed by DL28, nanobodies may achieve their high affinities by shape complementarity with antigen. Thus, the concave arc formed by CDR3 and the loop at the other end clamps onto the antigen. This type of interaction has also been observed in the case of nanobodies against the KDEL receptor ^38^, the κ-opioid receptor ^39^, the folate transporter ^40^, and the histo-blood group antigen BabB ^41^.

In the literature, increasing avidity generally improves potency, although the effect can vary from dozens to thousands of times ^10, 11^. Interestingly, the avidity effect for both DL4 was not apparent (**Table S1**), and that for DL28 was only obvious for the Beta variant (**Fig. 5C, Table S1**). Mechanistically, fusing with Fc may introduce additional steric hindrance to prevent RBD-ACE2 binding. It may also tether two S trimers to restrict their conformational changes should the two nanobody entities bind to different S. More commonly, avidity is known to increase potency by boosting apparent binding affinity by increasing local concentration and hence a faster *k*_on_ and a slower *k*_off_. In the case of DL4/DL28, the affinity may not be the limiting factor owing to their exceptional binding kinetics. This provides a possible reason for the lack of avidity effect. Despite this, the Fc fusion can increase the potency *in vivo* by extending the serum half-life of nanobodies from several minutes to several days ^10^ and thus should be still be useful for therapeutic reasons.

The fact that the DL4(3m) is more potent than DL4 is worth discussing. Thus, despite ultra-high affinity after multiple rounds of immunization, there is still space for rational design. Such practice may be applied to the existing antibodies although the effect of mutations on pharmacological behavior will have to be tested in the cases of therapeutic antibodies.

It was rationalized that the DL4 R50E/D mutants would gain at least some neutralization activity towards the Beta variant by restoring a salt bridge that was probably lost due to the E484K mutation. However, the results showed that mutant was as ineffective as the wildtype DL4 (**Table S2**). This may suggest that, apart from a sidechain replacement, subtle conformational changes also occur in this region and the changes are compatible with ACE2-binding but not for DL4. It is also possible that other mutations contained in the Beta variant, although not directly involved in the binding with DL4, helped escape the nanobody by allosteric effects. This highlights the challenges in the development of broadly effective neutralizing antibodies against SARS-CoV-2. Antibodies need high affinity to work best, but high affinity requires the epitope arranges in a precise three-dimensional shape. Mutation even remote from the epitope can distort the fine shape and render the antibodies ineffective. Targeting structurally rigid domains is key to develop broadly active antibodies.

Owing to their minute sizes, nanobodies may bind surfaces that are inaccessible for conventional antibodies. In the case of DL4 and DL28, they may be able to bind to the ‘down’-RBD given their minor clashes with the ‘closed’ conformation of S, in addition to binding with the ‘up’-RBD (**Fig. S1, S3**). On the other hand, the small size could mean that the destruction of S trimer by binding, as observed for several conventional antibodies, can be rare ^42, 43^.

Despite similar binding kinetics, DL28 showed less neutralizing activity (∼5 fold) compared to DL4. In addition, the cross-competition for ACE2-binding was complete by DL4 (**Fig. 3C**) but not by DL28 (**Fig. 7A**). We do not yet understand the structural reason for this. Possibly, ACE2 can interact weakly with RBD via the non-distorted part of the RBM. As also reported in the literature, RBM-targeting antibodies are generally more competent for neutralization, i.e., direct completion is generally more efficient for ACE2-blocking ^44, 45^. However, by targeting the more conserved RBD core region ^31^, antibodies that do not aim at the RBM may be less susceptible to escape mutants. Whether this is the case for DL28 remains to be investigated using replication-competent viruses.

Because DL28’s epitope only marginally overlaps with the RBM, DL28 may be able to bind RBD in the presence of other RBM-targeting nanobodies and human monoclonal antibodies. Such pairs will allow the development of biparatopic nanobodies to increase tolerance to escape mutants, and DL28’s ultra-high affinity could offer great advantages in such applications.

Although we did not test the neutralizing activity of DL4 and DL28 against the Delta strain, DL4 is expected to be a weak neutralizer because the critical Glu484’ for DL4-binding is, as in the case for the Beta variant, mutated, although the replacement is glutamine instead of lysine. For DL28, the impact of the Delta mutations is not very clear from the structural analysis. As shown in **Fig. S5**, Leu452’ is a part of a hydrophobic network comprised of Phe490’, Tyr351’, and Ile468’ from RBD, and Phe47, Tyr37, Ile50, and Trp104 from DL28. The Delta mutation L452R would weaken the hydrophobic interactions. Besides, the RBD Arg452’ may be repulsed by DL28 Arg45 in the vicinity. On the other hand, however, Arg452 may, depending on the sidechain conformations, form a cation-π interaction with Trp104, and/or form a salt bridge with Asp102. Thus, the exact effect will need to be tested in the future. In the case of weakened neutralizing activity, mutations to accommodate the Delta variant such as W104D can be designed and screened to restore neutralizing activity.

As revealed by the structure of the ACE2-RBD complex, ACE2 engages with RBD mainly through two structurally rigid α-helices (α_20-52_ and α_55-82_). By contrast, the counterpart in RBD is composed of loops lay on top of the core RBD region (**Fig. 7B, 7E**). This interaction model is perhaps suited for the RBD function. Thus, the structural flexibility of loops allows RBD to assume different, but functionally competent conformations by adjusting its backbone position while allowing escape mutants to evolve. However, the flexible feature also makes it prone to distortion, and the RBM distortion can have detrimental consequences for ACE2-binding, as demonstrated here by DL28. Our work suggests a previously unreported mechanism for SARS-CoV-2 neutralization which could be exploited for developing therapeutic nanobodies.

## CONCLUSIONS

We obtained two alpaca nanobodies that target RBD with ultra-high affinities and neutralize SARS-CoV-2 with high potencies. DL4 neutralizes SARS-CoV-2 by direct competition with ACE2 for RBD-binding, whereas DL28 distorts the ACE2-binding site and forces RBD to a conformation incompatible with receptor engagement. DL28 can neutralize the Alpha and Beta variants which are more infectious than the original SARS-CoV-2 strain. Our study provides two tight nanobodies for research and potential therapeutic applications and suggests an uncommon mechanism for SARS-CoV-2 neutralization.

## MATERIALS AND METHODS

### Protein expression and purification - Spike (S)

The polypeptide containing, from N- to C-terminus, residues Met1 – Gln1208 (without the C-terminal transmembrane helix, Uniprot P0DTC2) of the SARS-CoV-2 S with mutations K986P/V987P, a GSAS linker substituting the furin sites (Arg682- Arg685), a C-terminal T4 fibritin trimerization motif (GYIPEAPRDGQAYVRKDGEWVLLSTFL), a TEV protease cleavage site, a FLAG tag and a polyhistidine tag ^5^ was encoded in a pCDNA3.1 backbone vector and overexpressed in Expi293 cells by transient transfection using polyethylenimine (PEI). After 3.5 days of suspension culturing, the medium was harvested by filtration through a 0.22-μm membrane, and adjusted to contain 200 mM NaCl, 20 mM imidazole, 4 mM MgCl_2_, and 20 mM Tris-HCl pH 7.5. The filtrate was incubated with 3 mL of Ni-NTA beads at 4 °C for 2 h. The beads were loaded into a Bio-Rad gravity column, washed with 50 column volume (CV) of 20 mM imidazole, and subsequently eluted with 250 mM imidazole in 200 mM NaCl, 20 mM Tris-HCl pH 7.5. Fractions containing S were pooled, concentrated with a 100-kDa cut-off membrane concentrator, and further purified by gel filtration. S protein was quantified using a theoretical ε_280_ of 138,825 _M-1 cm-1._

### Protein expression and purification - RBD

The polypeptide containing, from N- to C-terminus, the honey bee melittin signal peptide (KFLVNVALVFMVVYISYIYAA), a Gly-Ser linker, residues 330-531 of the SARS-CoV-2 S, a Gly-Thr linker, the 3C protease site (LEVLFQGP), a Gly-Ser linker, the Avi tag (GLNDIFEAQKIEWHE), a Ser-Gly linker, and a deca-His tag was encoded in a pFastBac-backbone vector for overexpression in *Trichoplusia ni* High Five suspension cells. Cells at 2 × 10^6^ cells per milliliter were transfected with baculovirus generated using standard Bac-to-Bac procedures (Invitrogen) and the expression was allowed for 48-60 h at 27 °C in flasks. The medium from 1 L of culture was filtered using a 0.22-μm membrane and the filtrate was adjusted to contain 30 mM imidazole before incubating with 3.0 mL of Ni-Sepharose Excel (Cat 17-3712-03, GE Healthcare) beads for 2 h at 4 °C with mild agitation. The beads were loaded into a gravity column, washed with 10 CV of 20 mM imidazole, and eluted using 300 mM of imidazole in 150 mM NaCl, 20 mM Tris HCl pH 8.0. For site-specific biotinylation, the Avi-tagged RBD at 0.8 mg mL^-1^ was incubated with 5 mM ATP, 10 mM magnesium acetate, 43.5 μM biotin, 22 μg mL^-1^ home-purified BirA in a 3.2-mL reaction mix and incubated at 4 °C for 16 h. Biotinylated RBD was concentrated with a 10-kDa cut-off membrane to ∼3 mg mL^-1^ before loaded onto a Superdex Increase 200 10/300 GL column for gel filtration. Fractions containing the RBD were pooled, aliquoted, flash-frozen in liquid nitrogen, and stored at -80 °C before use.

For crystallization, RBD eluted from the Ni-NTA column was desalted using a desalting column, and digested with home-purified 3C protease to remove the C-terminal tags. The resulted tag-free RBD was mixed with nanobodies (see below) at a molar ratio of 1:1.3 and the mix was loaded onto a Superdex Increase 200 10/300 GL column for gel filtration. Fractions containing the complex were pooled, concentrated to 10 mg mL^-1^ for crystallization.

### Protein expression and purification - monovalent nanobodies in *Escherichia coli*

Monovalent nanobodies were expressed with a C-terminally Myc tag and a hexahistidine tag in *E. coli* MC1061 cells. Briefly, cells carrying nanobody-encoding pSb-init plasmids ^46^ were grown in Terrific Broth (TB, 0.17 M KH_2_PO_4_ and 0.72 M K_2_HPO_4_, 1.2 %(w/v) tryptone, 2.4 %(w/v) yeast extract, 0.5% (v/v) glycerol) supplemented with 25 mg L^-1^ chloramphenicol at 37 °C with shaking at 200 rpm. When cell density reached an OD_600_ of 0.5 (∼ 2 h), the shaker was set to 22 °C and the cells were allowed to grow for another 1.5 h before added with 0.02% (w/v) arabinose for induction for 17 h. Cells were harvested by centrifugation and lysed by osmotic shock as follows. Briefly, cells from 1 L of culture were resuspended in 20 mL of TES-high Buffer (0.5 M sucrose, 0.5 mM EDTA, and 0.2 M Tris-HCl pH 8.0) and incubated at 4 °C for 30 min. Dehydrated cells were then abruptly rehydrated using 40 mL of ice-cold MilliQ H_2_O at 4 °C for 1 h to release periplasmic protein. The periplasmic extract was collected by centrifugation at 20,000 g at 4 °C for 30 min. The supernatant was adjusted to have 150 mM of NaCl, 2 mM of MgCl_2_, and 20 mM of imidazole before incubated with Ni-NTA beads that had been pre-equilibrated with 20 mM of imidazole, 150 mM NaCl, and 20 mM Tris HCl pH 8.0. After batch-binding for 2 h, the Ni-NTA beads were washed using 30 mM imidazole, before eluted using 300 mM imidazole, 150 mM NaCl, and 20 mM Tris HCl pH 8.0. Nanobodies were quantified using their theoretical molar extinction coefficient calculated based on the contents of aromatic residues.

### Protein expression and purification - divalent nanobodies in mammalian cells

Nanobodies with a C-terminal Fc fusion and an N-terminal leader peptide (MEFGLSWVFLVALLRGV) were transiently expressed in Expi293 suspension cells. Briefly, cells at 2.5 × 10^6^ cells per milliliter were transfected with a mix of plasmids and PEI. Valproic acid was included at 2 mM to increase expression. After 65 h at 37 °C, the medium was harvested by centrifugation at 100×*g* and filtration. The filtrate was incubated with rProtein A beads (Cat SA012005, SmartLifesciences, China) for batch binding at 4 °C for 3 h. The beads were packed into a gravity column, washed using 20 CV of PBS buffer before eluted using 0.1 M glycine pH 3.0. The elution was immediately neutralized with 1 M Tris HCl pH 8.0. The buffer was then exchanged to PBS on a Bio-Rad desalt column.

Nanobody mutants in this study were all generated on the Fc-fusion constructs using standard PCR-based site-directed mutagenesis protocols. DNA sequences were verified by sequencing and the mutants were expressed and purified the same way as their wild-type proteins.

### Protein expression and purification - monoclonal antibodies

DNA encoding the heavy and light chain variable regions of the monoclonal antibodies (mAbs) REGN10933, CV30, and CB6 were synthesized and separately Gibson-assembled into a pDEC vector which features the human IgG backbone. For each mAb, two plasmids (1.4 mg L^-1^ for light chain, 0.6 mg L^-1^ for heavy chain), were co-transfected into Expi293 cells using polyethylenimine at a cell density of 2 × 10^6^ per milliliter for transient expression. Valproic acid was added to a final concentration of 2 mM to increase the expression level. The medium containing secreted IgG-mAbs was collected by centrifugation 48-60 h post-transfection, filtered using a 0.22-μm membrane, and incubated with Protein A affinity beads for 2 h at 4 °C. The beads were transferred into a gravity column and washed with 20 CV of PBS buffer before eluted with 0.1 M glycine pH 3.0. The acidic eluent was rapidly neutralized by 1 M Tris-HCl pH 8.0. NaCl was adjusted to a final concentration of 0.15 M. The purified mAbs were buffer-exchanged into PBS using a desalting column (Bio-Rad). mAbs were quantified using their theoretical molar extinction coefficients that are calculated based on the contents of aromatic residues and with absorbance at 280 nM measured using a Nanodrop machine.

### Alpaca immunization and antibody titer determination

Purified RBD (1 mL at 2 mg mL^-1^) was mixed with an equal volume of the Gerbu adjuvant (Cat. 3111) by vortexing. The resulted emulsion was injected by the subcutaneous route at ten sites near the bow lymph node in the neck base of an adult female alpaca (3-years old). The immunization process was repeated 3 times (a total of 4 rounds) with 4 days between each injection.

To determine the antibody titer, 3 mL of blood samples before and after each injection were collected. After 2 h at room temperature (RT, 20-25 °C), the clotted sample was centrifuged at 3,000 g for 5 min at RT to collect sera in the supernatant. Wells of 96-well plates (Maxisorp, Nunc Thermo Fisher Scientific) were coated overnight at 4 °C with 100 μL of 2 μg mL^-1^ biotinylated RBD in TBS (150mM NaCl, 20mM Tris, pH8.0) and blocked with 0.5% bovine serum albumin (BSA) in TBS. After washing five times with TBS, serially diluted alpaca sera were added and incubated for 1 h. After washing, the bound nanobody was detected by HRP-conjugated Goat anti-Alpaca IgG (Cat. S001P, NBbiolab) using Tetramethylbenzidine (TMB) (Merck, Cat.T2885) as a substrate for HRP. ELISA test of sera showed an antibody titer of ∼1 × 10^6^ after four rounds of immunization compared with the pre-immunization sample.

### Phage display library construction and panning

Eighty milliliters of blood were collected from the immunized alpaca in EDTA-coated tubes. The tubes were inverted twice to inhibit coagulation. The peripheral blood lymphocytes were isolated using Ficoll Plus (density of 1.077 g mL-1) according to the manufacturer’s instructions. Isolated lymphocytes were used for mRNA isolation with RNAsio Plus (TaKara). Reverse transcription was performed using mRNA and a commercial kit (Vazyme Cat. R312-01). Polymerase chain reaction (PCR) was carried out with 50 ng of cDNA and the primer pair CALL001 (5’- GTCCTGGCTGCTCTTCTACAAGG-3’) and CALL002 (5’- GGTACGTGCTGTTGAACTGTTCC-3’) using the PCR Master Mix (Cat. 10149ES01, YEASEN Biotech, Shanghai, China). The PCR product was loaded onto a 1.5 %(w/v) agarose gel and the 700-bp band was excised. The purified PCR product was used for a second round of PCR using the prime pair VHH-BspQI-F (5’-ATAT*GC TCTTC*AAGTCAGGTGCAGCTGCAGGAGTCTGGRGGAGG-3’) and VHH- BspQI-R (5’-TATA*GCTCTTC*CTGCCGAGGAGACGGTGACCTGGGT-3’) which anneals to the framework 1 and framework 4 region of nanobodies, respectively. The primers contained a recognition site (italic) for the type IIs restriction enzyme *BspQ*I for cloning purposes. The PCR product was purified using a FastPure kit (Vazyme Cat. DC301).

One microgram of the PCR product and 10 μg of the pDX_init vector ^46^ were digested separately with 50 units of *BspQ*I (Cat. R0712L, New England Biolabs) for 1.5 h at 50 °C before heat inactivation at 80 °C for 10 min. The digested DNA were gel-purified and 0.3 μg of the PCR product were mixed with 1.2 μg of vector and 10 units of T4 ligase in ligation buffer (Cat. B110041, Sangon Biotech, Shanghai, China) for 1.5 h. The mixture was transformed into *Escherichia coli* SS320 cells by electroporation in a 2-mm cuvette using a Gene Pulser Xcell (Bio-Rad) with a setting of 2,400 volts, 25 μF, and 750 Ω.

Cells were grown in 225 mL of 2-YT broth (1.0 %(w/v) yeast extract, 1.6 %(w/v) tryptone, 0.5 %(w/v) NaCl, pH 7.0) supplemented with 200 μg mL^-1^ ampicillin and 2 %(w/v) glucose in a 37-°C shaking incubator at 220 rpm. To 10 mL of the overnight culture, 27 μL of the M13KO7 helper phage at 10^12^ plaque-forming units mL^-1^ were added. After brief mixing, the mixture was incubated at 37 °C for 30 min. The cells were collected by centrifugation at 3,200×*g* for 10 min, resuspended in 2-YT broth supplemented with 200 μg mL^-1^ ampicillin and 25 μg mL^-1^ kanamycin, and placed in a shaker incubator at 37 °C with 160 rpm.

After 16 h of culture, the medium from 50 mL of culture was collected by centrifugation at 3,200×*g* for 30 min at 4 °C. The supernatant (40 mL) was transferred to a fresh Falcon tube. Phage particles were precipitated by incubating the supernatant with 10 mL of 20 %(w/v) PEG 6,000 and 2.5 M NaCl for 30 min on ice. Precipitated phage particles were collected by centrifugation at 3,200×*g* for 30 min at 4 °C before resuspended in 1 mL of PBS buffer. After centrifugation at 20,000×*g* for 5 min, the supernatant was transferred into a fresh 1.5-mL tube and the procedure was repeated once.

The first round was performed in a Nunc Maxisorp 96-well immunoplate. The plate was first coated with 67 nM neutravidin (Cat. 31000, Thermo Fisher Scientific) overnight at 4 °C, followed by blocking with TBS buffer supplemented with 0.5 %(w/v) BSA for 30 min. Phage particles (4.9 mL) were incubated with 50 nM biotinylated RBD, added to the neutravidin-coated wells, washed, and released from the plate by tryptic digestion (10 min at RT) with 0.25 mg mL^-1^ trypsin in the buffer containing 150 mM NaCl and 20 mM Tris-HCl pH 7.4. After being treated with the trypsin inhibitor AEBSF, the selected phage particles were amplified in *E. coli*, and the second solution panning was performed as the first round except that the plate was replaced with 12 μL of MyOne Streptavidin C1 beads (Cat. 65001, Invitrogen). The bound phage particles were challenged with 5 μM non-biotinylated RBD to compete off binders with fast off-rates. The third round of panning was performed the same as the second round except that the RBD concentration was at 5 nM. The particles were eluted, and the phagemid was sub-cloned into pSb_init vector by fragment-exchange (FX) cloning and transformed into *E. coli* MC1061 cells for periplasmic expression and screening.

### Enzyme-linked immunosorbent assay (ELISA) – nanobody selection

Single colonies carrying pSb-init plasmids above were grown at 37 °C for 5 h in a shaking incubator at 300 rpm before 1:20 seeded into 1 mL of fresh TB supplemented with 25 μg mL^-1^ chloramphenicol. Cells were induced with 0.02% (w/v) arabinose at 22 °C for 17 h before collected by centrifugation at 3,000 g for 30 min. Cell pellets were resuspended in TES Buffer (20 % (w/v) sucrose, 0.5 mM EDTA, 0.5 μg/mL lysozyme, 50 mM Tris-HCl pH 8.0) and incubated for 30 min at room temperature (RT, 20-25 °C). The lysate was added with 0.9 mL of TBS (150 mM NaCl, 20 mM Tris-HCl pH 7.4) supplemented with 1 mM MgCl_2_. The mix was centrifuged at 3,000 g for 30 min at 4 °C and the supernatant containing nanobodies was used for ELISA as follows.

Wells of a Maxi-Sorp plate (Cat. 442404, Thermo Fisher) was coated with Protein A at 4 °C for 16 h. The plate was then blocked by 0.5 %(w/v) BSA in TBS buffer for 30 min at RT and washed three times using TBS before incubated with anti-Myc antibodies at 1:2,000 dilution in TBS-BSA-T buffer (TBS supplemented with 0.5 %(w/v) BSA and 0.05 %(v/v) Tween 20) for 20 min at RT. The plate was then washed three times with TBST (TBS supplemented with 0.05% Tween 20) to remove excess antibodies. The wells were incubated with the Myc-tagged nanobodies prepared above for 20 min at RT. After washing three times with TBST, the wells were incubated with 50 nM of biotinylated RBD or MBP (the maltose-binding protein, as a control) for 20 min at RT. The wells were again washed three times with TBST before incubated with streptavidin-conjugated with horseradish peroxidase (HRP) (1:5,000, Cat S2438, Sigma). After 30 min, the plate was washed three times with TBST. ELISA signal (absorbance at 650 nm) was developed by incubating the wells with 100 μL of developing reagents (51 mM Na_2_HPO_4_, 24 mM citric acid, 0.006 %(v/v) H_2_O_2_, 0.1 mg mL^-1^ 3,3’,5,5’-tetramethylbenzidine) at RT.

### Fluorescence-detection size exclusion chromatography (FSEC) – nanobody selection

FSEC analysis of RBD-binding by nanobodies was performed as previously described ^10^. Biotinylated RBD was incubated with streptavidin (Cat 16955, AAT Bioquest) that was chemically labeled by fluorescein. The fluorescent complex (500 nM) was mixed with the cell lysate containing unpurified nanobodies and the mixture was applied onto an analytic gel filtration column (Cat 9F16206, Sepax) connected to an HPLC system equipped with a fluorescence detector (RF-20A, Shimadzu) for FSEC analysis. The FSEC profile was monitored by fluorescence at the excitation/emission pair of 482/508 nm and compared to that incubated with a control MBP-nanobody for peak shift.

### Biolayer interferometry for S-nanobody binding and competitive binding

The binding kinetics was measured by a bio-layer interferometry (BLI) assay using an Octet RED96 system (ForteBio). A streptavidin-coated SA sensor (Cat. 18-5019, Sartorius) was coated with 5 μg mL^-1^ biotinylated nanobodies for approximately 1 min. The sensor was equilibrated in a nanobody-free buffer for ∼30 s, before bathing in solutions containing various concentrations (association) of Spike (analytes) for 120 s (DL4) or 360 s (DL28).

For competition between ACE2 and nanobodies, biotinylated RBD (2 μg mL^-1^) was immobilized on an SA sensor by incubating with the sensor in the BLI Buffer (0.005 %(v/v) Tween 20, 1 × phosphate-buffered saline) at 30 °C. The RBD-loaded sensor was saturated in 100 nM of nanobodies for 6-15 min. The sensor was then bathed in nanobody solutions with or without 100 nM of ACE2 (Cat 10108-H08B). The association of ACE2 was monitored for 360 s. As a control, the ACE2-RBD binding profile was recorded using the same procedure as above but in the absence of nanobodies.

For simultaneous binding of DL28 and monoclonal antibodies (REGN10933, CV30, CB6, all in the IgG form) with RBD, the biotinylated RBD was coated as mentioned above. The sensor was then equilibrated with 100 nM of DL28 (monovalent). The sensor was then bathed in DL28-containing solution (100 nM) with or without 100 nM of individual mAbs for BLI recording. As controls, the RBD-mAb binding was recorded in the same manner without DL28.

Data were fitted for a 1:1 stoichiometry for *K*_D_, *k*_on_, and *k*_off_ calculations using the built-in software Data Analysis 10.0.

### Crystallization

Crystallization trials were set up in a two-well sitting-drop plate with 70 μL of reservoir solution, and 1 μL each of the protein solution and the precipitant solution. The plates were incubated at 16 °C for crystal growth. The precipitant solution for the DL4-RBD complex contained 25 %(w/v) polyethylene glycol 4,000, 0.2 M ammonium sulfate, and 0.1 M sodium acetate pH 4.6. The crystallization condition for DL28-RBD contained 20 %(w/v) polyethylene glycol 3,350, and 0.2 M potassium phosphate dibasic. Cryo protection was achieved by adding 20 %(v/v) glycerol in the respective precipitant condition. Crystals were harvested using a MitGen loop, and flash-cooled in liquid nitrogen before X-ray diffraction data collection.

### X-ray data collection and structure determination

X-ray diffraction data were collected at beamline BL18U1 at Shanghai Synchrotron Radiation Facility with a 50 × 50 μm beam on a Pilatus detector at a distance of 300 – 500 mm, with oscillation of 0.5° and a wavelength of 0.97915 Å. Data were integrated using XDS ^47, 48^, and scaled and merged using Aimless ^49^. The structure was solved by molecular replacement using Phaser ^50^ with the RBD structure (PDB 6M0J) ^31^ and a nanobody structure (PDB 5M13) ^46^ as the search model. The model was built with 2*F*_o_-*F*_c_ maps in Coot ^51^, and refined using Phenix ^52^. Structures were visualized using PyMol.

### Data availability

The structure factors and coordinates are available through the protein data bank (PDB) under accession codes 7F5G (DL4-RBD) and 7F5H (DL28-RBD).

### Neutralization assay using SARS-CoV-2 pseudoviruses

Retroviral pseudotyped particles were generated by co-transfection of HEK293T cells using polyethylenimine with the expression vectors encoding the various viral envelope glycoproteins, the Murine leukemia virus core/packaging components (MLV Gag-Pol), and a retroviral transfer vector harboring the gene encoding the green fluorescent protein (GFP). The S Protein expressed by phCMV-SARS-CoV-2 has been truncated to remove 19 amino acid residues at the C-terminal. Supernatants that contained pseudotyped particles were harvested 48 h post-transfection and filtered through a 0.45-μm membrane before neutralizing assays.

VeroE6-hACE2 cells (10^4^ cells/well) were seeded into a 48-well plate and infected 24 h later with 100 μL of virus supernatant in a final volume of 150 μL. Nanobodies were pre-incubated with the pseudotype samples for 1 h at 37 °C before cell/virus co-incubation. After 6 h of co-incubation, the supernatants were removed and the cells were incubated in the medium for 72 h at 37 °C. GFP expression was determined by fluorescence-activated flow cytometry analysis (FACS). The infectivity of pseudotyped particles incubated with nanobodies was compared with the infectivity using pseudotyped particles and Dulbecco’s modified Eagle’s medium-2% fetal calf serum only and normalized to 100%.

Average and standard deviation (s.d., n = 3) were plotted for the IC_50_ experiments except for Fc-DL4(3m) in Fig. 4C and 5B which report data from two independent experiments.

SARS-CoV-2 pseudotypes for the Alpha (B1.1.7) and Beta (B1.351) variants were generated by incorporating the corresponding Spike mutations into the phCMV-SARS-CoV-2 plasmid. Desired mutations were verified by DNA sequencing.

### Animal experiment and ethics

The alpaca immunization procedures were conducted in conformity with the institutional guidance for the care and use of laboratory animals, and the protocols were approved by the Institutional Committee of Ethics and Research of the Central Laboratory at Xinyang Agricultural and Forestry University.

## CONFLICT OF INTERESTS

Patent applications for potential nanobody therapy for the treatment of COVID-19 have been filed for DL4 and DL28.

## AUTHOR CONTRIBUTION

T.L. and Y.L. selected, purified, and characterized nanobodies under the supervision of D.Li. B.Z. and S.H. performed neutralizing assays under the supervision of D.La.. Y.Z. and S.W. immunized alpaca and constructed phage-display library. Y.Z. performed crystallization and collected X-ray diffraction data under the supervision of J.T.. A.G. and S.B. constructed Spike mutants and helped establish neutralization assays.

J.B. helped with protein engineering. D.Li and Z.L. processed X-ray diffraction data and solved structures. D.Li analyzed structure, designed mutants, and wrote the manuscript with input from T.L. and D.La..

## ACKNOWLEDGEMENT

We thank the staff scientists at the SSRF-BL18U1 beamline at National Facility for Protein Science (Shanghai) for technical support, Dr. Peiling Li from Xinyang Agricultural and Forestry University for assistance in the alpaca immunization, and Ms. Yuchen Li for sketching Fig. 1A. This work has been supported by the Strategic Priority Research Program of CAS (XDB37020204), Key Program of CAS Frontier Science (QYZDB-SSW-SMC037), CAS Facility-based Open Research Program, the National Natural Science Foundation of China (31870726, D.Li; 31870153, D.La.), National Key R&D Program of China (2020YFC0845900, D.La.), CAS president’s international fellowship initiative (2020VBA0023, D.La.), Natural Science Foundation of Shanghai (20ZR1466700, D.Li), Shanghai Municipal Science and Technology Major Project (20431900402, D.La.).

## Supplementary Information

**Table S1.**
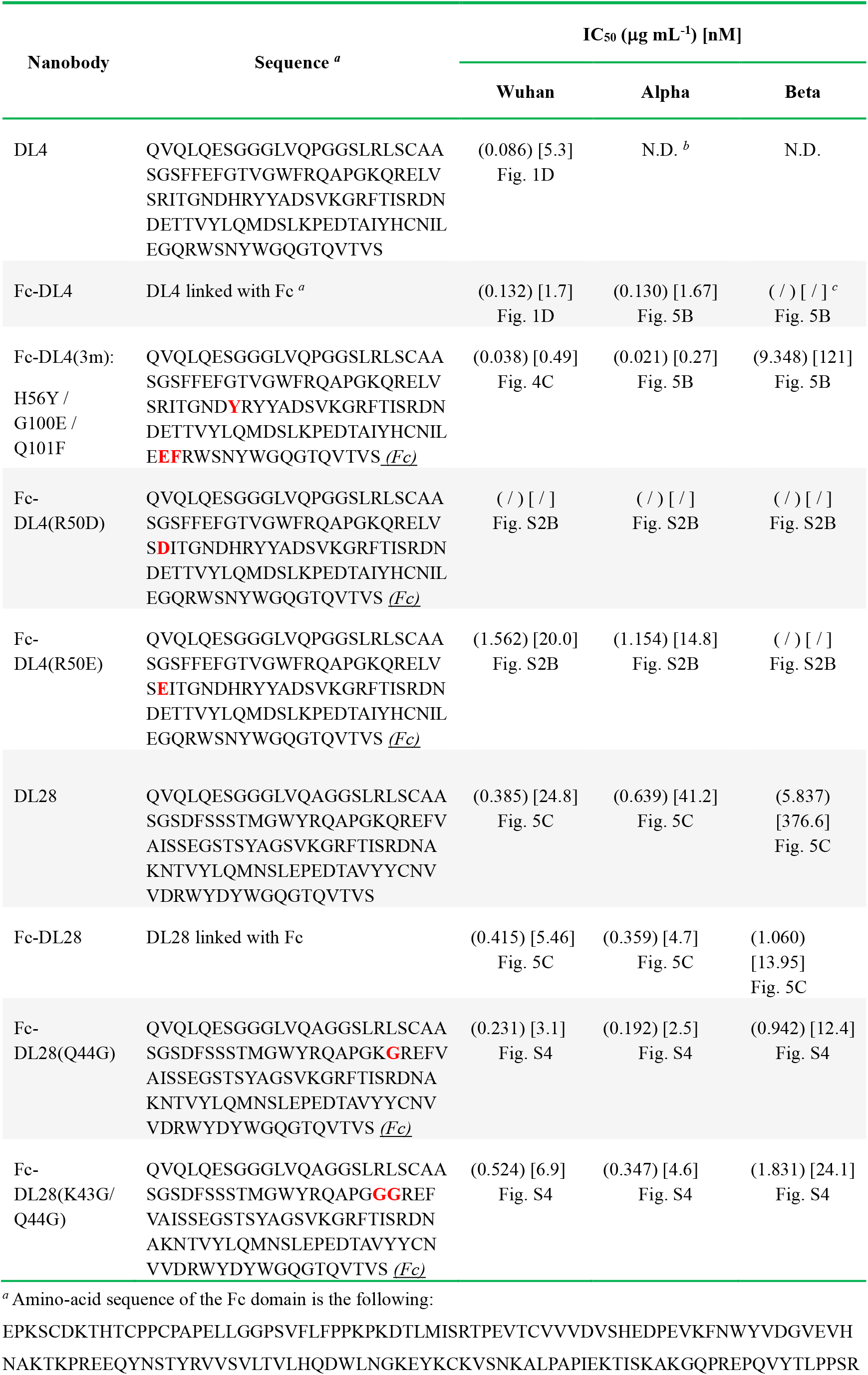
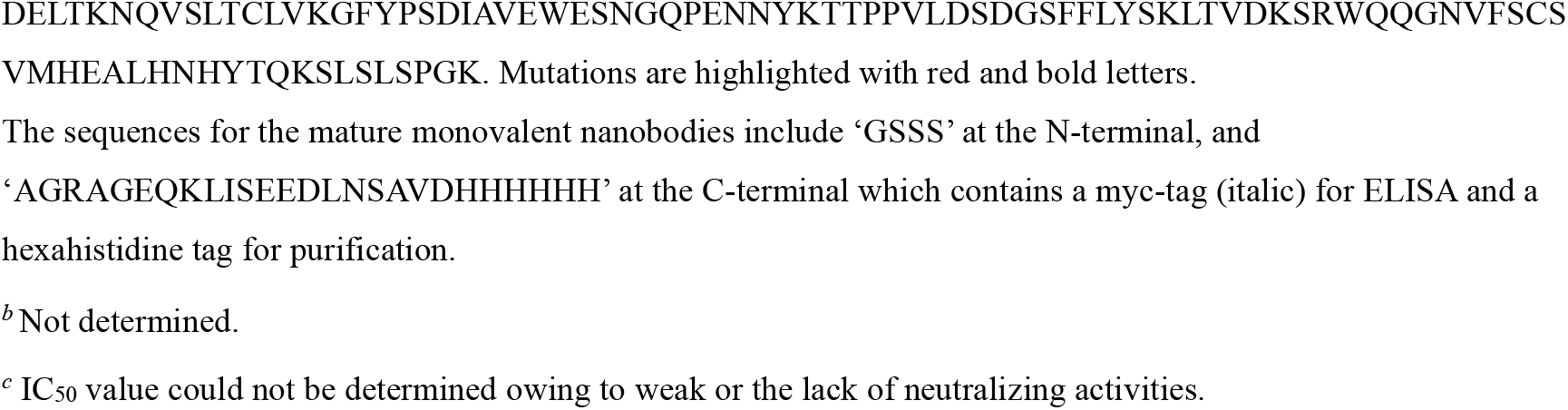
Summary of nanobody.

**Table S2.**
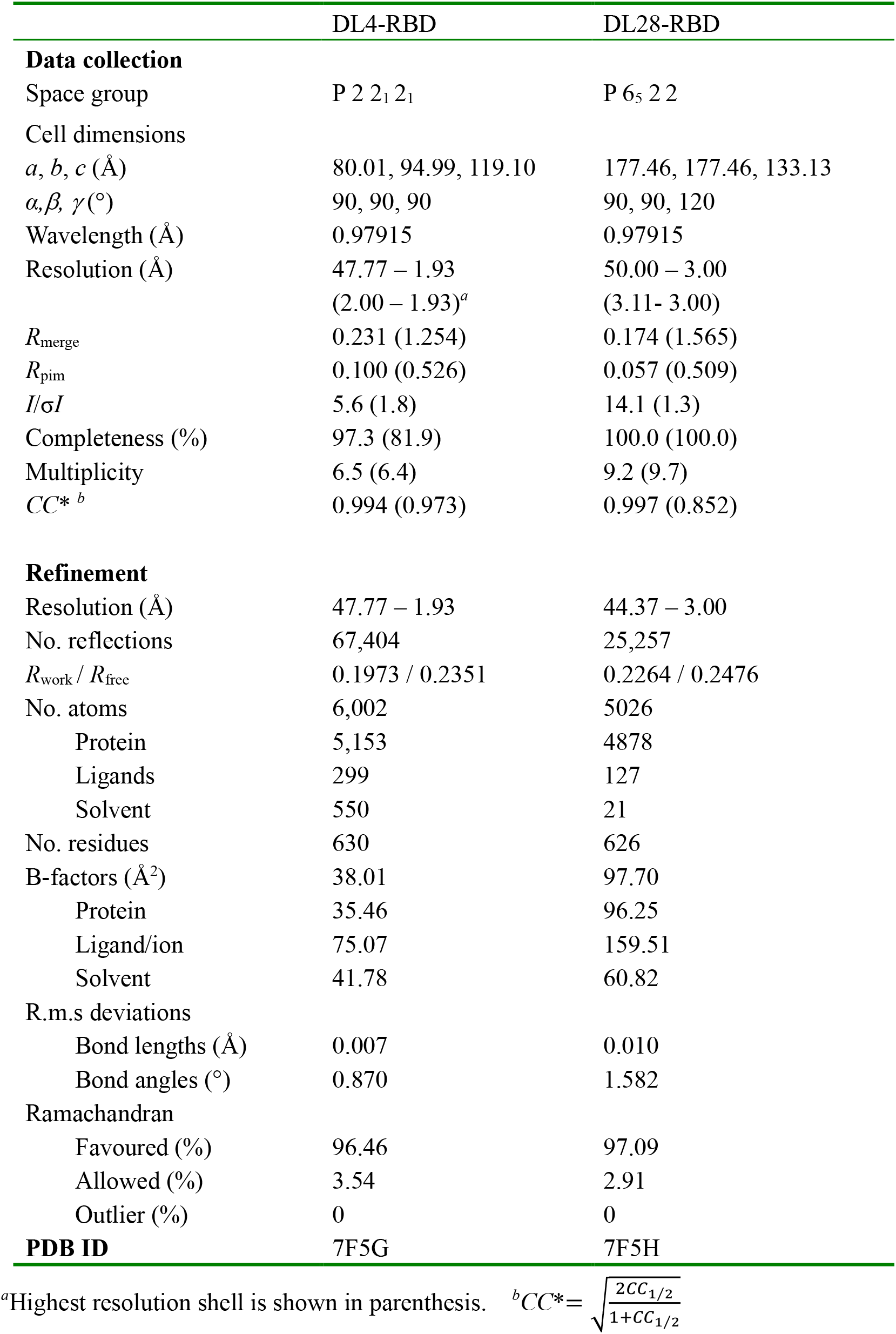
Data collection and refinement statistics.

**Fig. S1.**
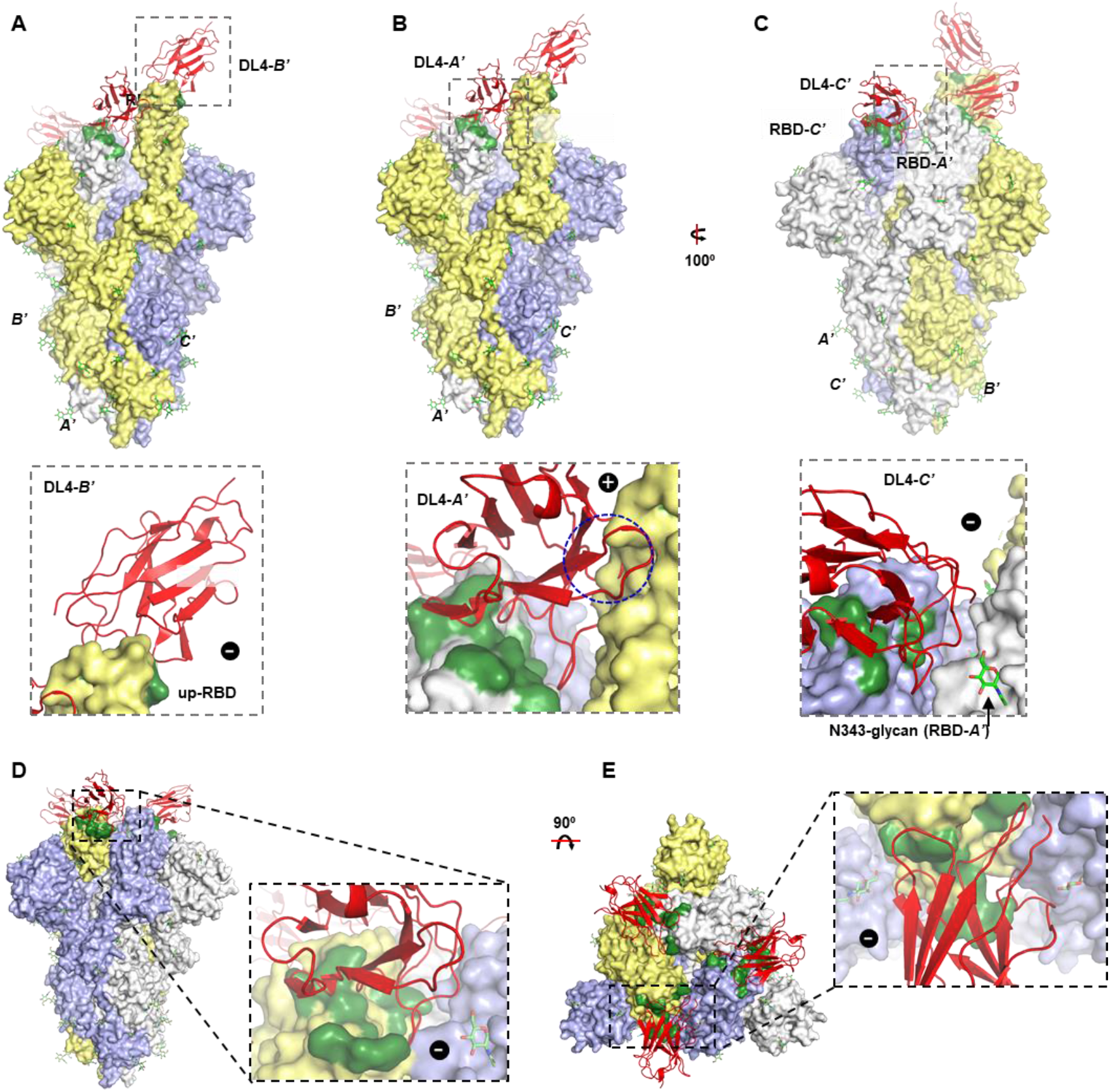
Docking the DL4 to the SARS-CoV-2 Spike. (**A-C**) DL4 is aligned to the open conformation of Spike (S) (PDB ID 6VYB) ^1^ on its ‘up’-RBD (**A**) and two ‘down’-RBDs (**B** and **C**). Three subunits are labeled with *A’*, *B’,* and *C’*. DL4 docked to the subunits are marked with the individual *A’/B’/C’* subunits. The expanded views are shown below each panel. ‘+’ denote clashes (blue circle) and ‘-’ denotes minor or the lack of clashes. (**D, E**) Side view (**D**) and top view (**E**, viewed from the membrane normal) of S in the closed conformation (PDB ID 6VXX) ^1^ with docked DL4. Because the three subunits in the closed conformation are identical, the interactions are only shown for one subunit.

**Fig. S2.**
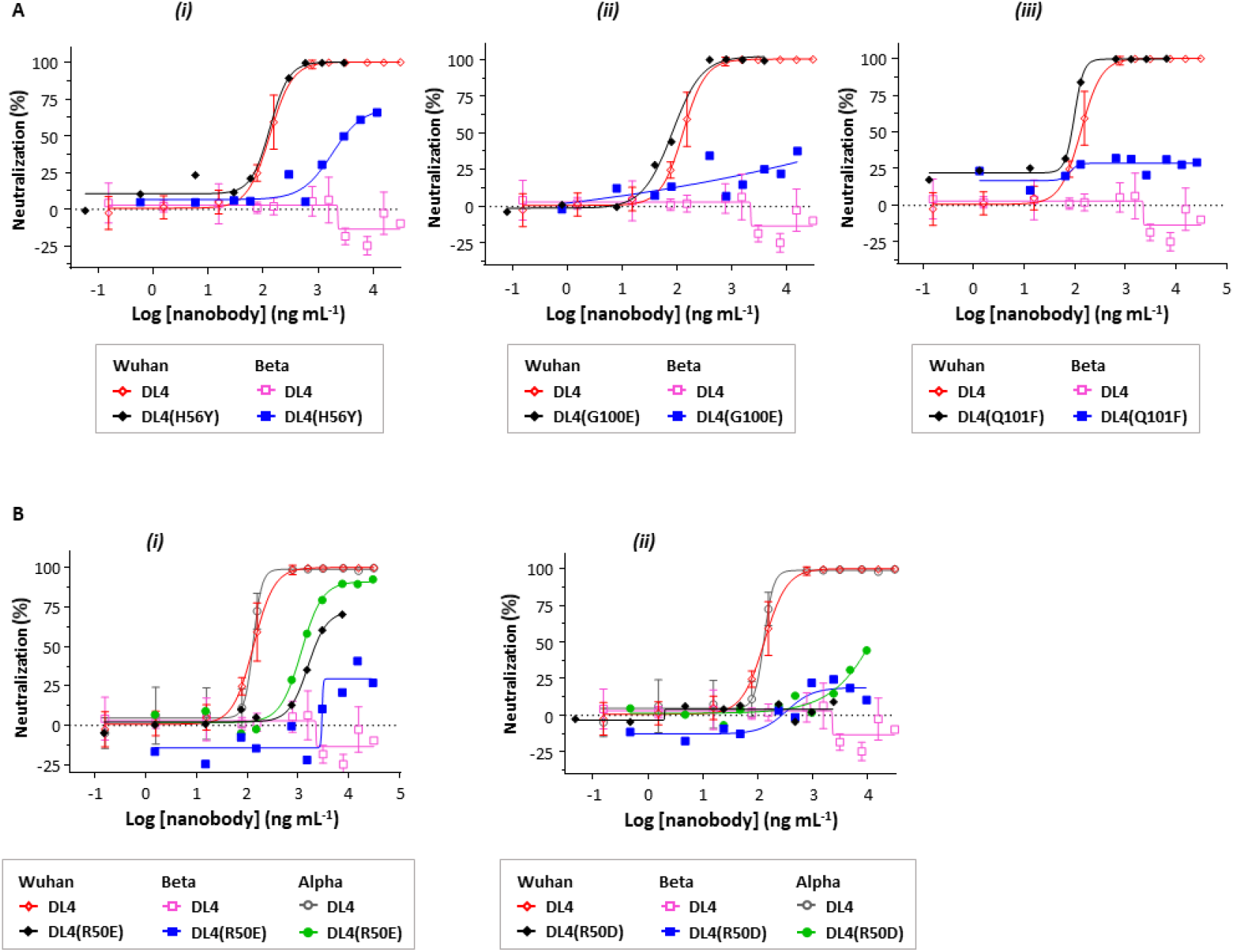
Neutralizing assay of DL4 mutants. (**A**) Neutralization assay of three gain-of-function single-point mutants H56Y (***i***), G100E (***ii***), and Q101F (***iii***). (**B**) Neutralizing assay for DL4(R50E) (***i***) and DL4(R50D) (***ii***). The data for the wild-type DL4 are from Fig. 1C. Fc-nanobodies were used for neutralization assays. Data for mutants are from a single experiment.

**Fig. S3.**
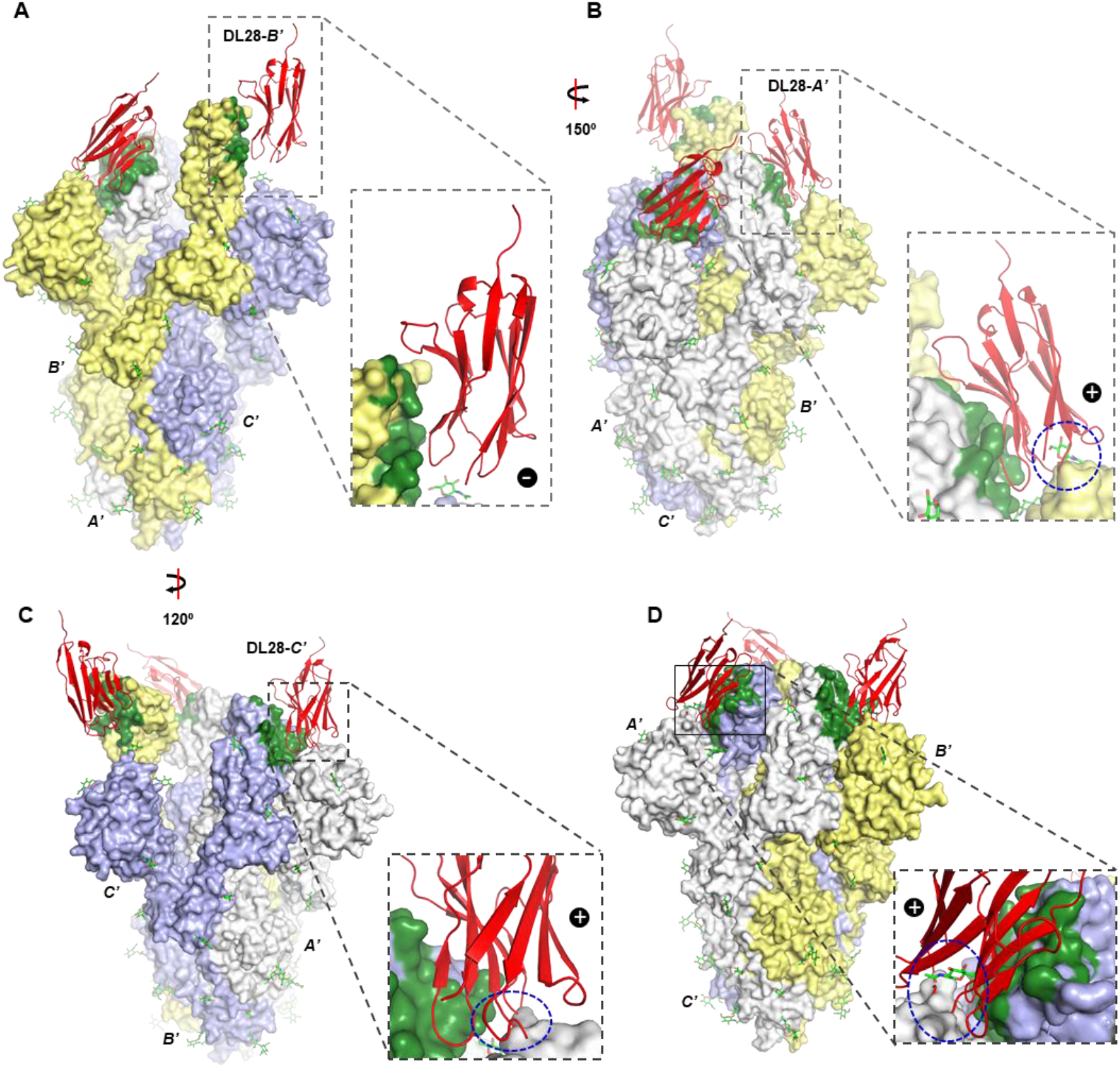
Docking the DL28 to the SARS-CoV-2 Spike. (**A-C**) DL28 is aligned to the open conformation of Spike (S) (PDB ID 6VYB) ^1^ on its ‘up’-RBD (**A**) and two ‘down’-RBDs (**B** and **C**). Three subunits are labeled with *A’*, *B’,* and *C’*. DL28 docked to the subunits are marked with the individual *A’/B’/C’* subunits. ‘+’ denote clashes (blue circle) and ‘-’ denotes lack of clashes. (**D**) DL28 clashes with the closed conformation of S (PDB ID 6VXX) ^1^. Because the three subunits in the closed conformation are identical, the interactions are only shown for one subunit.

**Fig. S4.**
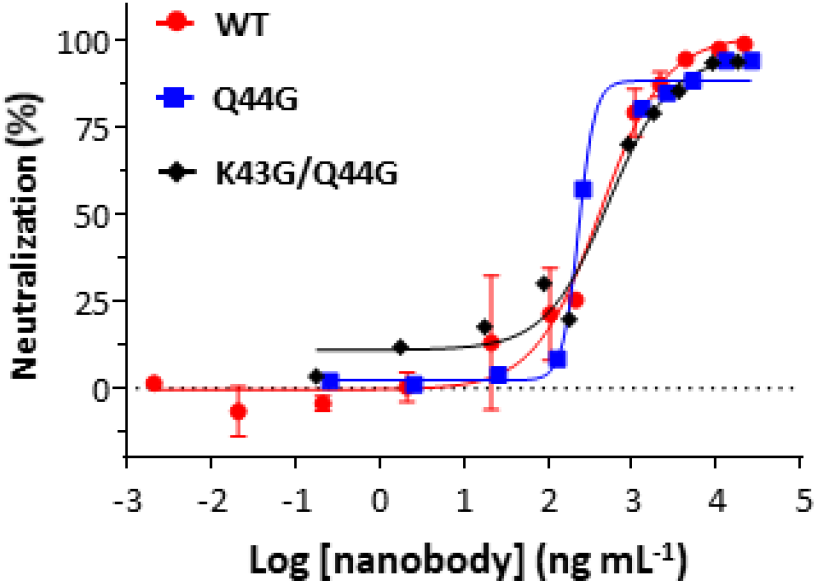
Steric hindrance by Gln44 does not contribute to neutralization. Neutralization assays for Q44G and K43G/Q44G using the original SARS-CoV-2 Wuhan strain. The data for DL28 are from Fig. 5C for comparison reasons. Fc- nanobodies were used in the assay. Data for the two mutants are the average of two independent experiments.

**Fig. S5.**
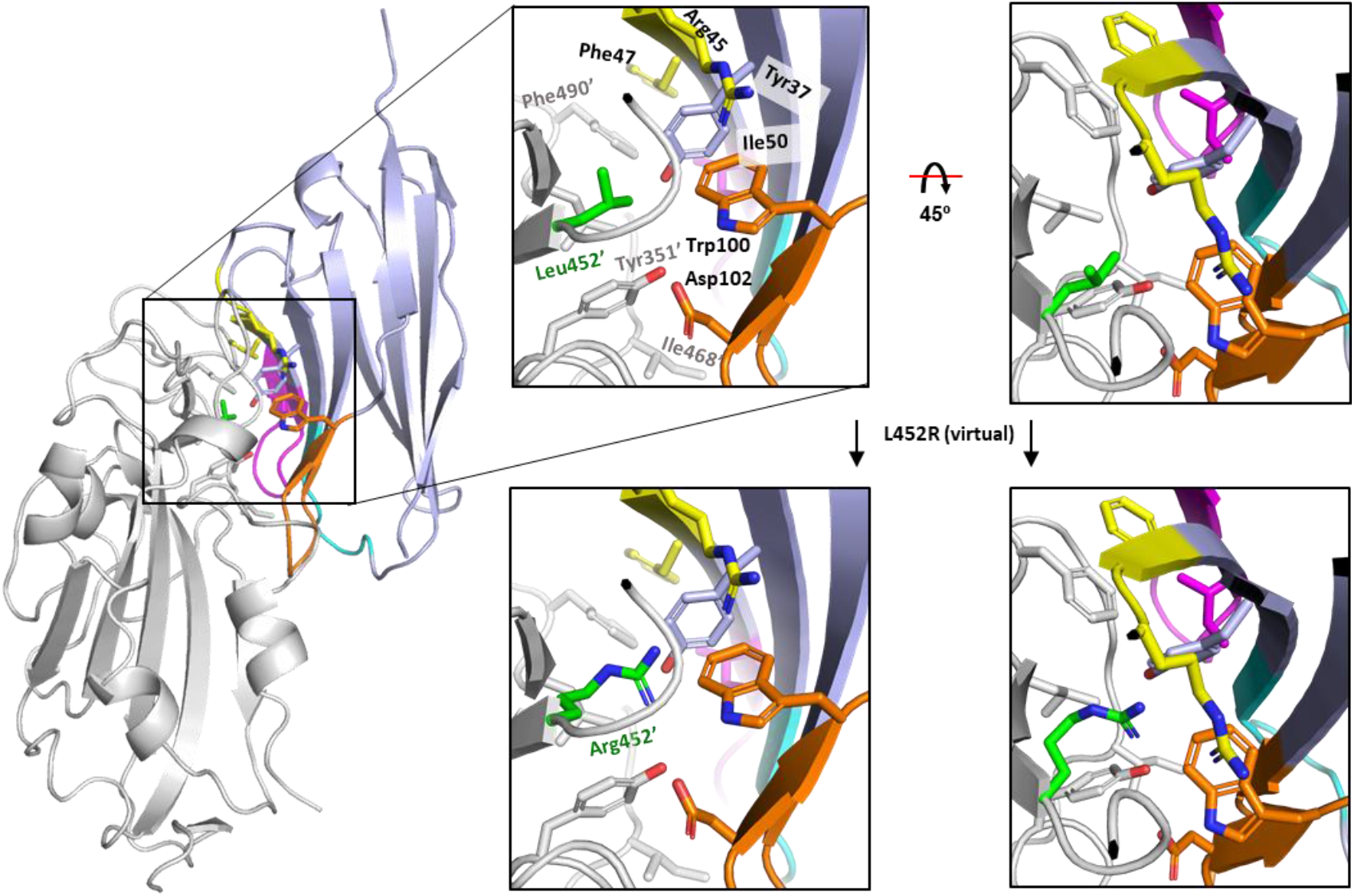
Virtual mutation of RBD L452R and the possible consequences. Leu452 was virtually mutated to arginine. The surrounding residues on both RBD (white) and DL28 (light blue with CDR1 in Cyan, CDR2 in magenta, CDR3 in orange, and RBD-interacting framework residues in yellow) are shown as sticks. RBD residues are labeled with a prime. Leu452’ is part of a hydrophobic network formed by the shown residues. In the Delta variant, Arg452’ may become incompatible with the hydrophobic microenvironment. On the other hand, however, Arg452’ may, depending on the sidechain conformations, form a salt bridge with DL28 Asp102.

## REFERENCES

1. Walls, A.C., et al. Structure, Function, and Antigenicity of the SARS-CoV-2 Spike Glycoprotein. Cell 181, 281–292.e6 (2020).

2. Wrapp, D. et al. Cryo-EM structure of the 2019-nCoV spike in the prefusion conformation. Science 367, 1260–1263 (2020).

3. Shang, J. et al. Cell entry mechanisms of SARS-CoV-2. Proceedings of the National Academy of Sciences 117, 11727–11734 (2020).

4. Hoffmann, M. et al. SARS-CoV-2 Cell Entry Depends on ACE2 and TMPRSS2 and Is Blocked by a Clinically Proven Protease Inhibitor. Cell 181, 271–280.e8 (2020).

5. Zhang, C. et al. Development and structural basis of a two-MAb cocktail for treating SARS-CoV- 2 infections. Nature Communications 12, 264 (2021).

6. Henderson, R. et al. Controlling the SARS-CoV-2 spike glycoprotein conformation. Nature Structural & Molecular Biology 27, 925–933 (2020).

7. Barnes, C.O. et al. SARS-CoV-2 neutralizing antibody structures inform therapeutic strategies. Nature 588, 682–687 (2020).

8. Muyldermans, S. Nanobodies: natural single-domain antibodies. Annu Rev Biochem 82, 775–97 (2013).

9. Yao, H. et al. A high-affinity RBD-targeting nanobody improves fusion partner’s potency against SARS-CoV-2. PLOS Pathogens 17, e1009328 (2021).

10. Li, T. et al. Potent synthetic nanobodies against SARS-CoV-2 and molecular basis for neutralization. bioRxiv, 2020.06.09.143438 (2020).

11. Schoof, M. et al. An ultrapotent synthetic nanobody neutralizes SARS-CoV-2 by stabilizing inactive Spike. Science 370, 1473–1479 (2020).

12. Koenig, P.-A. et al. Structure-guided multivalent nanobodies block SARS-CoV-2 infection and suppress mutational escape. Science 371, eabe6230 (2021).

13. Nambulli, S. et al. Inhalable Nanobody (PiN-21) prevents and treats SARS-CoV-2 infections in Syrian hamsters at ultra-low doses. bioRxiv, 2021.02.23.432569 (2021).

14. Pymm, P. et al. Nanobody cocktails potently neutralize SARS-CoV-2 D614G N501Y variant and protect mice. Proceedings of the National Academy of Sciences 118, e2101918118 (2021).

15. Walter, J.D. et al. Synthetic nanobodies targeting the SARS-CoV-2 receptor-binding domain. bioRxiv, 2020.04.16.045419 (2020).

16. Custódio, T.F. et al. Selection, biophysical and structural analysis of synthetic nanobodies that effectively neutralize SARS-CoV-2. Nature Communications 11, 5588 (2020).

17. Hanke, L. et al. An alpaca nanobody neutralizes SARS-CoV-2 by blocking receptor interaction. Nature Communications 11, 4420 (2020).

18. Huo, J. et al. Neutralizing nanobodies bind SARS-CoV-2 spike RBD and block interaction with ACE2. Nature Structural & Molecular Biology 27, 846–854 (2020).

19. Chi, X. et al. Humanized single domain antibodies neutralize SARS-CoV-2 by targeting the spike receptor binding domain. Nature Communications 11, 4528 (2020).

20. Esparza, T.J., Martin, N.P., Anderson, G.P., Goldman, E.R. & Brody, D.L. High affinity nanobodies block SARS-CoV-2 spike receptor binding domain interaction with human angiotensin converting enzyme. Scientific Reports 10, 22370 (2020).

21. Xiang, Y. et al. Versatile and multivalent nanobodies efficiently neutralize SARS-CoV-2. Science 370, 1479–1484 (2020).

22. Starr, T.N., Greaney, A.J., Dingens, A.S. & Bloom, J.D. Complete map of SARS-CoV-2 RBD mutations that escape the monoclonal antibody LY-CoV555 and its cocktail with LY-CoV016. Cell Reports Medicine 2, 100255 (2021).

23. Harvey, W.T. et al. SARS-CoV-2 variants, spike mutations and immune escape. Nature Reviews Microbiology 19, 409–424 (2021).

24. Weisblum, Y. et al. Escape from neutralizing antibodies by SARS-CoV-2 spike protein variants. eLife 9, e61312 (2020).

25. Liu, Z. et al. Identification of SARS-CoV-2 spike mutations that attenuate monoclonal and serum antibody neutralization. Cell Host & Microbe 29, 477–488.e4 (2021).

26. Wang, P. et al. Antibody resistance of SARS-CoV-2 variants B.1.351 and B.1.1.7. Nature 593, 130–135 (2021).

27. Hoffmann, M. et al. SARS-CoV-2 variants B.1.351 and P.1 escape from neutralizing antibodies. Cell 184, 2384–2393.e12 (2021).

28. Bolze, A. et al. Rapid displacement of SARS-CoV-2 variant B.1.1.7 by B.1.617.2 and P.1 in the United States. medRxiv, 2021.06.20.21259195 (2021).

29. Starr, T.N. et al. Prospective mapping of viral mutations that escape antibodies used to treat COVID-19. Science 371, 850–854 (2021).

30. Krissinel, E. & Henrick, K. Inference of Macromolecular Assemblies from Crystalline State. Journal of Molecular Biology 372, 774–797 (2007).

31. Lan, J. et al. Structure of the SARS-CoV-2 spike receptor-binding domain bound to the ACE2 receptor. Nature 581, 215–220 (2020).

32. Shang, J. et al. Structural basis of receptor recognition by SARS-CoV-2. Nature 581, 221–224 (2020).

33. Wrapp, D. et al. Structural Basis for Potent Neutralization of Betacoronaviruses by Single-Domain Camelid Antibodies. Cell 181, 1004–1015.e15 (2020).

34. Hansen, J. et al. Studies in humanized mice and convalescent humans yield a SARS-CoV-2 antibody cocktail. Science 369, 1010–1014 (2020).

35. Hurlburt, N.K. et al. Structural basis for potent neutralization of SARS-CoV-2 and role of antibody affinity maturation. Nature Communications 11, 5413 (2020).

36. Shi, R. et al. A human neutralizing antibody targets the receptor-binding site of SARS-CoV-2. Nature 584, 120–124 (2020).

37. Ye, G., et al. The Development of a Novel Nanobody Therapeutic for SARS-CoV-2. bioRxiv : the preprint server for biology, 2020.11.17.386532 (2020).

38. Bräuer, P. et al. Structural basis for pH-dependent retrieval of ER proteins from the Golgi by the KDEL receptor. Science 363, 1103–1107 (2019).

39. Che, T. et al. Structure of the Nanobody-Stabilized Active State of the Kappa Opioid Receptor. Cell 172, 55–67.e15 (2018).

40. Parker, J.L. et al. Structural basis of antifolate recognition and transport by PCFT. Nature (2021).

41. Moonens, K. et al. Structural Insights into Polymorphic ABO Glycan Binding by *Helicobacter pylori*. Cell Host & Microbe 19, 55–66 (2016).

42. Huo, J. et al. Neutralization of SARS-CoV-2 by Destruction of the Prefusion Spike. Cell Host & Microbe 28, 445–454.e6 (2020).

43. Zhou, D. et al. Structural basis for the neutralization of SARS-CoV-2 by an antibody from a convalescent patient. Nature Structural & Molecular Biology 27, 950–958 (2020).

44. Dejnirattisai, W. et al. The antigenic anatomy of SARS-CoV-2 receptor binding domain. Cell 184, 2183–2200.e22 (2021).

45. Piccoli, L. et al. Mapping Neutralizing and Immunodominant Sites on the SARS-CoV-2 Spike Receptor-Binding Domain by Structure-Guided High-Resolution Serology. Cell 183, 1024–1042.e21 (2020).

46. Zimmermann, I. et al. Synthetic single domain antibodies for the conformational trapping of membrane proteins. eLife 7, e34317 (2018).

47. Kabsch, W. XDS. Acta Crystallographica Section D 66, 125–132 (2010).

48. Kabsch, W. Integration, scaling, space-group assignment and post-refinement. Acta Crystallographica Section D 66, 133–144 (2010).

49. Evans, P.R. & Murshudov, G.N. How good are my data and what is the resolution? Acta Crystallogr D Biol Crystallogr 69, 1204–14 (2013).

50. McCoy, A.J., Storoni, L.C. & Read, R.J. Simple algorithm for a maximum-likelihood SAD function. Acta Crystallographica Section D 60, 1220–1228 (2004).

51. Emsley, P. & Cowtan, K. Coot: model-building tools for molecular graphics. Acta Crystallographica Section D 60, 2126–2132 (2004).

52. Adams, P.D. et al. PHENIX: a comprehensive Python-based system for macromolecular structure solution. Acta Crystallographica Section D 66, 213–221 (2010).

